# Dissecting the cellular specificity of smoking effects and reconstructing lineages in the human airway epithelium

**DOI:** 10.1101/612747

**Authors:** Katherine C. Goldfarbmuren, Nathan D. Jackson, Satria P. Sajuthi, Nathan Dyjack, Katie S. Li, Cydney L. Rios, Elizabeth G. Plender, Michael T. Montgomery, Jamie L. Everman, Eszter K. Vladar, Max A. Seibold

**Affiliations:** Center for Genes, Environment, and Health, National Jewish Health, Denver, CO, 80206 USA; Department of Pediatrics, National Jewish Health, Denver, CO, 80206 USA; Division of Pulmonary Sciences and Critical Care Medicine, University of Colorado-AMC, Aurora, CO, 80045 USA; Department of Cell and Developmental Biology, University of Colorado-AMC, Aurora, CO, 80045 USA

## Abstract

Cigarette smoke first interacts with the lung through the cellularly diverse airway epithelium and goes on to drive development of most chronic lung diseases. Here, through single cell RNA-sequencing analysis of the tracheal epithelium from smokers and nonsmokers, we generated a comprehensive atlas of epithelial cell types and states, connected these into lineages, and defined cell-specific responses to smoking. Our analysis inferred multi-state lineages that develop into surface mucus secretory and ciliated cells and contrasted these to the unique lineage and specialization of submucosal gland (SMG) cells. Our analysis also suggests a lineage relationship between tuft, pulmonary neuroendocrine, and the newly discovered CFTR-rich ionocyte cells. Our smoking analysis found that all cell types, including protected stem and SMG populations, are affected by smoking, through both pan-epithelial smoking response networks and hundreds of cell type-specific response genes, redefining the penetrance and cellular specificity of smoking effects on the human airway epithelium.

## Introduction

The human airway epithelium is a complex, cellularly diverse tissue that plays a critical role in respiratory health by facilitating air transport, barrier function, mucociliary clearance, and the regulation of lung immune responses. These airway functions are accomplished through interactions among a functionally diverse set of both common (ciliated, mucus secretory, basal stem) and rare cell types (pulmonary neuroendocrine, tuft, ionocyte) which compose the airway surface epithelium. This remarkable cell diversity derives from the basal airway stem cell by way of multiple branching lineages^1,2^, yet, the nature of these lineages, their transcriptional regulation and the functional heterogeneity to which they lead, remain incompletely defined in humans. Equally important to airway function, if even more poorly understood on both a molecular and cellular level, is the epithelium of the airway submucosal glands (SMG), a network that is contiguous with the surface epithelium and a critical source of airway mucus and defensive secretions.

Gene expression and histological studies of the airway epithelium have demonstrated that both molecular dysfunction and cellular imbalance due to shifting cell composition in the epithelium are common features of most chronic lung diseases, including asthma^3^ and chronic obstructive pulmonary disease^4^ (COPD). This cellular remodeling is largely mediated by interaction of the epithelium with inhaled agents such as air pollution, allergens, and cigarette smoke, which are risk factors for these diseases. Among these exposures, cigarette smoke is the most detrimental and, as the primary driver of COPD^5^ and a common trigger of asthma exacerbations^6^, constitutes the leading cause of preventable death in the U.S.^7^ Smoking is known to induce mucus metaplasia^8^ and gene expression studies based on bulk RNA-sequencing have established the dramatic influence of this exposure on airway epithelial gene expression^9–12^. However, these bulk expression changes are a composite of all cell type expression changes, frequency shifts, and emergent metaplastic cell states, making it impossible to determine the precise cellular and molecular changes induced by smoke exposure using this type of expression data.

Here, we use single cell RNA-sequencing (scRNA-seq) to define the transcriptional cell types and states of the tracheal airway epithelium in smokers and nonsmokers, infer the lineage relationship among these cells, and determine the influence of cigarette smoke on individual surface and SMG airway epithelial cell types with single cell resolution.

## Results

### Smoking induces both shared and unique gene expression responses across diverse airway epithelial cell types

To interrogate the cellular diversity of the human tracheal epithelium, we enzymatically dissociated tracheal specimens from seven donors and subjected these cells to scRNA-seq (Figure 1a). These donors included never-smokers, light smokers and heavy smokers (Supplementary Table S1), allowing us to evaluate the transcriptional effects of smoking habit on each epithelial cell type. Shared nearest neighbor (SNN) clustering of expression profiles from 13,840 epithelial cells identified eight broad cell clusters, each containing the full range of donors and smoking habits (Figure 1b, Supplementary Figure S1ab). Between 200-1500 differentially expressed genes (DEGs) distinguished these clusters from one another (Figure 1c, Supplementary Figure S1c).

**Figure 1:**
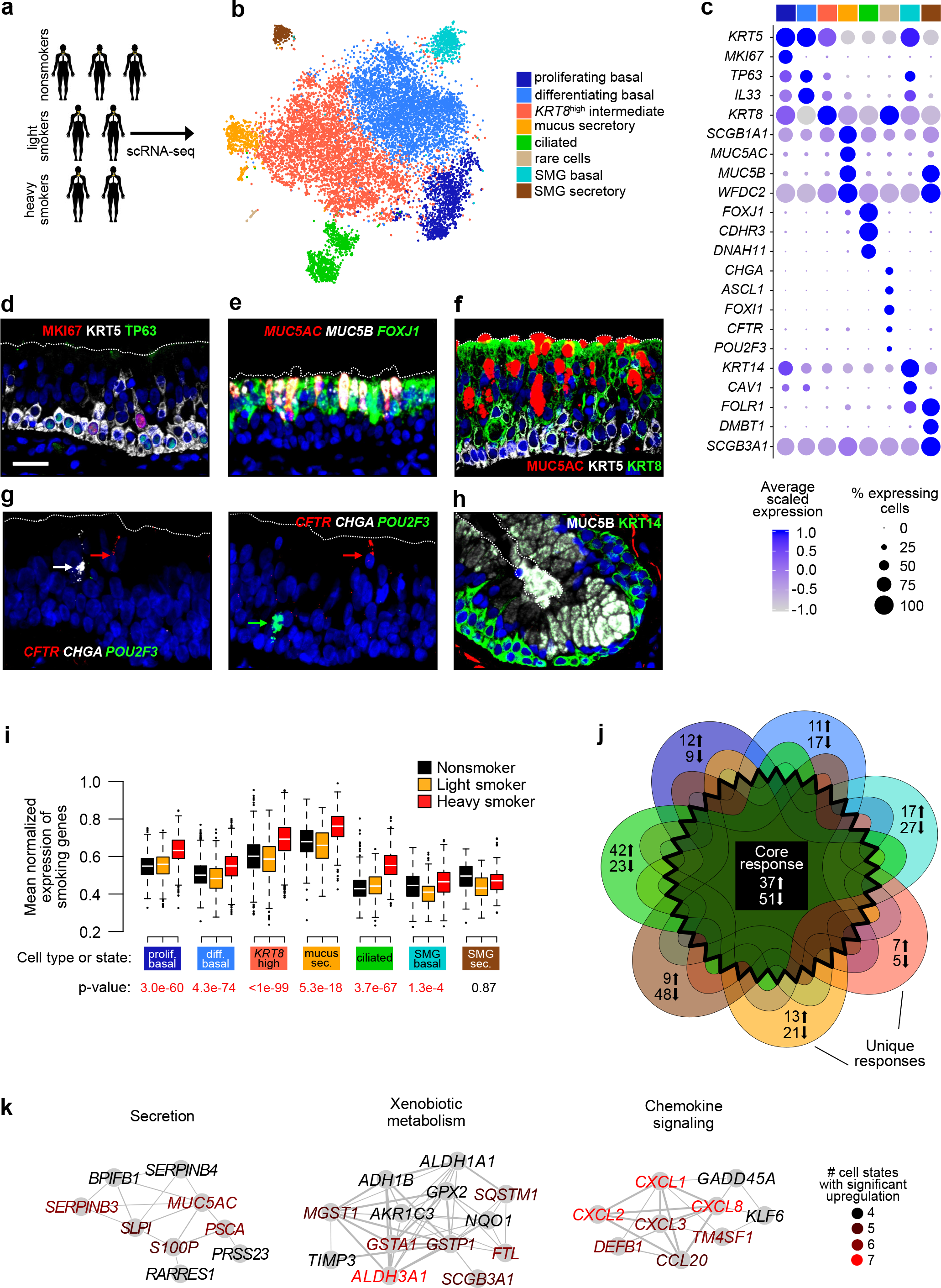
Shared and cell type-specific gene expression responses to smoking across diverse airway epithelial cell types. **a.** Schematic for studying human tracheal epithelium by scRNA-seq. **b.** tSNE visualization of cells in the human trachea depicts unsupervised clusters defining broad cell types present. **c.** Dot plots highlight known and novel markers distinguishing broad cell categories based on average expression level (color) and ubiquity (size). Colored bar corresponds to the cell types/states in **b**. **d.** Immunofluorescence (IF) labeling of human tracheal sections shows MKI67 (red) and TP63 (green) in a subset of KRT5^high^ (white) basal cells. Dotted line denotes the approximate apical surface of epithelium. Scale bar = 25 μm. DAPI labeling of nuclei (blue). **e.** Fluorescence *in situ* hybridization (FISH) co-localizes *MUC5B* (white) and
*MUC5AC* (red) mRNA to non-ciliated cells. *FOXJ1* (ciliated marker, green). **f.** IF labeling localizes KRT8 (green) to mid-upper epithelium, MUC5AC (red), KRT5 (white). **g.** FISH distinguishes rare cell mRNA markers: *Left*, PNECs (*CHGA*, white) and ionocytes (*CFTR*, red); *Right*, ionocytes and tuft-like cells (*POU2F3*, green). **h.** IF labeling localizes MUC5B (white) and KRT14 (green) to distinct cells in the SMGs. **i.** Published bulk RNA-seq upregulated smoking genes^9^ are upregulated in most heavy smoker cell populations. Box plots show distributions of mean normalized expression, p-values are based on one-sided t-tests comparing means for heavy and non-smokers. Rare cells were excluded due to small cell numbers. Box plots for downregulated genes from the same published study are in Supplementary Figure S1f. **j.** Venn diagram summarizes core and unique smoking responses across seven broad cell populations (colors match those in **b**), with number of up and downregulated genes unique to populations given in the tips and number of core genes affected in ≥ four populations in the center. Note degree of overlap in the diagram is not proportional to gene overlap for readability. Detailed percentages in Supplementary Figure S1g. **k.** Protein-protein interaction (PPI) networks show shared function across core smoking upregulated genes. Redness indicates number of cell populations where a gene was significantly upregulated (FDR < 0.05). See also Supplementary Figure S1, Supplementary Tables S1-S2.

Assignment of clusters to cell types or states was accomplished by examining expression of known epithelial cell type markers^13,14^ (Figure 1c). High *KRT5* expression identified two basal cell populations, distinguished by the presence/absence of proliferation markers (e.g. *MKI67*, Supplementary Figure S1c). Proliferative heterogeneity within *KRT5*^high^ basal cells was confirmed by immunofluorescence (IF) labeling in tracheal tissue sections (Figure 1d). Expression of the two major airway gel-forming mucins^15^, *MUC5AC* and *MUC5B*, identified the mucus secretory population via fluorescence *in situ* hybridization (FISH) (Figure 1ce). The ciliated cell population was identified through expression of ciliogenesis and ciliary function markers, including *FOXJ1*^16^ and *DNAH11*^17^ (Figure 1ce). We also identified a cluster characterized by high *KRT8* expression. Consistent with KRT8 being a differentiating epithelial cell marker^18^, KRT8^high^ cells localized to the mid-to-upper epithelium, above KRT5^high^ basal cells and often reaching the airway surface by IF (Figure 1f). Gene expression across *KRT8*^high^ cells was highly heterogeneous, with a wide range of expression for both basal (*KRT5*, *TP63*) and early secretory cell (*SCGB1A1*, *WFDC2*) markers. The smallest cluster contained cells expressing markers diagnostic for pulmonary neuroendocrine cells or PNECs^19^ (*CHGA*, *ASCL1*), ionocytes^20,21^ (*FOXI1*, *CFTR*) and tuft cells^22^ (*POU2F3*). Although these cells comprised less than 1% of the total epithelium, FISH of tracheal tissue identified cells expressing each of these markers (Figure 1g).

In addition to surface epithelial populations, two clusters highly expressed known glandular genes^23^ (Figure 1c, Supplementary Figure S1d), suggesting our digest isolated SMG epithelial cells. One of these SMG clusters highly expressed basal markers (e.g. *KRT5*, *KRT14*), while the other exhibited a mucus secretory cell character, including high *MUC5B* expression. IF labeling of tracheal tissue using these markers allowed visualization of SMG secretory and basal cells (Figure 1h, Supplementary Figure S1e).

Each identified population contained cells from all three smoking groups (Supplementary Figure S1b) and thus, we investigated the transcriptomic effects of smoking habit in each of the cell populations independently. We first examined smoking effects of genes previously reported to be differentially expressed between current and never-smokers using bulk RNA-seq data from bronchial airway epithelial brushings^9^. For both the upregulated and downregulated gene lists reported, a significant shift in mean expression between heavy smokers and nonsmokers was observed in six of seven populations that matched the direction of effect in the bulk data (Figure 1i, Supplementary Figure S1f). These effects, however, were not consistently observed in light smokers (Supplementary Figure S1f), leading us to focus further investigation of smoking effects on heavy smoker vs. nonsmoker cells.

Transcriptome-wide single-gene differential expression analysis identified over 150 DEGs between heavy and nonsmokers in each cell population (Supplementary Figure S1g, Supplementary Table S2). Importantly, 7%-87% of the smoking DEGs for each population were unique to that population, revealing a previously unappreciated cell type-specific aspect to the smoking response, discussed below (Figure 1j, Supplementary Figure S1g). Additionally, we identified a “core” response to heavy smoking, encompassing genes consistently up-or downregulated in at least four populations. Among these, *MUC5AC* was notably upregulated in six of seven (non-rare) cell types, while protein-protein interaction (PPI) network analysis of upregulated core genes revealed a pan-epithelial induction of nine interacting secretion-related genes with heavy smoking (Figure 1k). The upregulated core response also included genes enriched for xenobiotic metabolism and chemokine signaling (Figure 1k), suggesting that known airway responses to smoking, like toxin metabolism and macrophage recruitment^10,24^, are a joint effort conducted across epithelial cell types. The downregulated core response largely involved deactivation of immune function, such as the complement system, which helps clear microbes and damaged cells, and secretoglobin 1A1 (*SCGB1A1*) production, important for airway defense^25^ (Supplementary Figure S1i). Notably, the downregulated core response contained multiple HLA type I and II genes (Supplementary Figure S1i), possibly signaling an underappreciated role for antigen presentation in the epithelium, which is suppressed by smoking.

### Secretory cells form a continuous lineage that culminates in mucus secretory cells

Airway secretory cells canonically include club cells, which produce SCGB1A1-laden defensive secretions, and mucus secretory cells, which varyingly express the major gel-forming mucins (MUC5AC and MUC5B). Although inflammatory stimuli have been shown to induce conversion of club cells into mucus cells in mice^26^, the lineage relationship between these cells in the homeostatic human airway is unclear. Moreover, while NOTCH signaling is a likely mediator of secretory cell fate in the differentiating airway^27,28^, and the transcription factor (TF), SPDEF^29,30^, specifically drives inflammation-induced mucus metaplasia, little else is known regarding regulation of human secretory cell development. To investigate this area, we reconstructed the human secretory cell lineage using pseudotime trajectory analysis^31^ of the mucus secretory cells and *KRT8*^high^ populations, which contained cells with both an intermediate basal/secretory profile and club-like cells. This analysis aligned most cells along a single lineage (Supplementary Figure S2a) in which basal-like cells transitioned into mucus secretory cells through expression of three successive gene modules (Figure 2a). These modules included TFs and signaling molecules that may drive their expression (Figure 2b). The first of these modules (secretory preparation) was highly enriched for genes involved in ATP production and protein translation elongation, likely reflecting necessary preparation for the high energy demands of secretory protein production (Figure 2a). Secretory preparation genes were enriched for NOTCH signaling and included the *NOTCH3* receptor, as well as potential novel TF regulators (*BTF3*, *KLF3*) (Figure 2ab).

**Figure 2:**
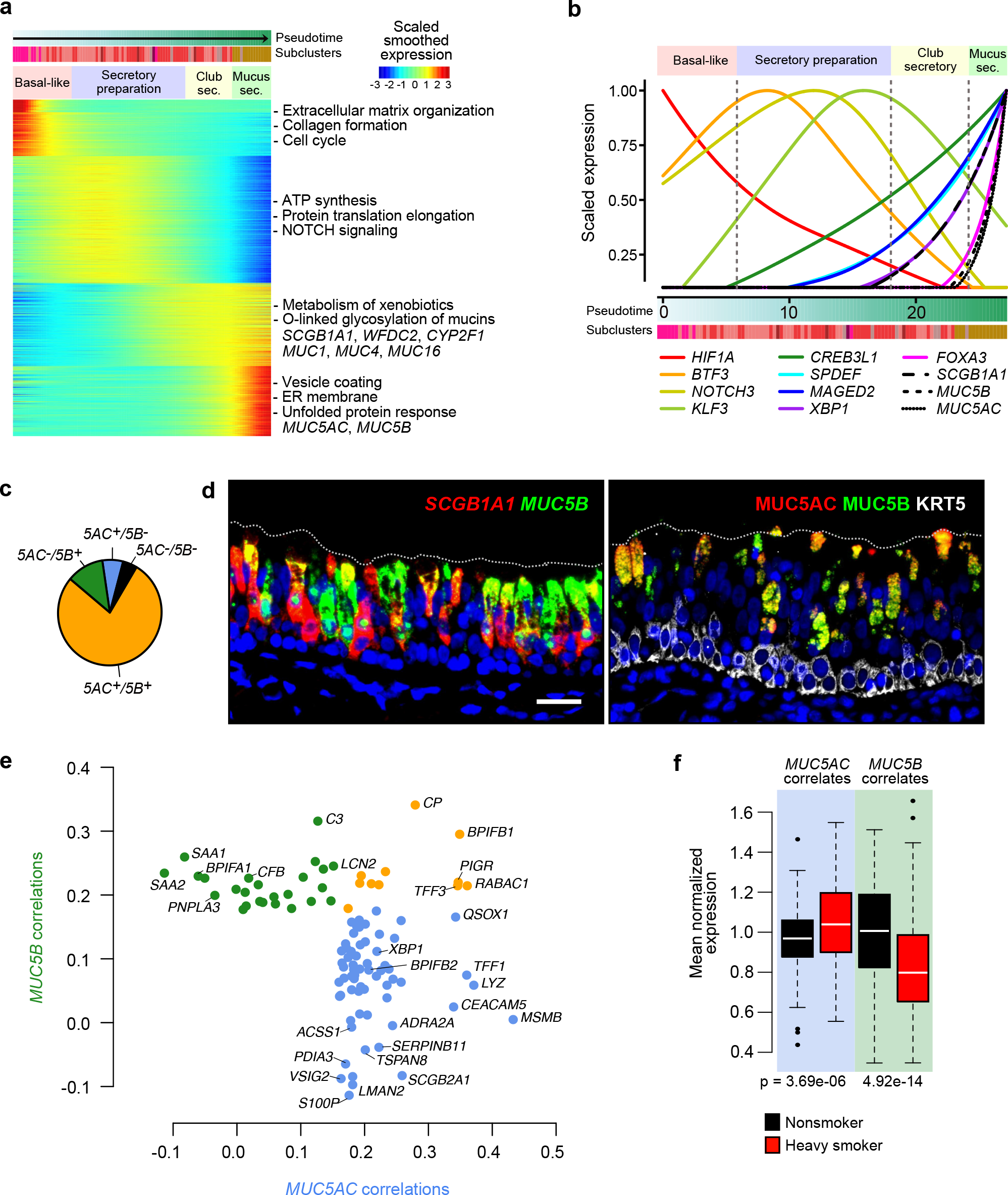
*In vivo* secretory cells form a continuous lineage and exhibit *MUC5AC*-correlated smoking effects. **a.** Heat map of smoothed expression across a Monocle-inferred lineage trajectory shows transitions in transcriptional programs that underlie differentiation in the *in vivo* human airway epithelium, from basal-like pre-secretory (*KRT8*^high^) cells into mucus secretory cells. Select genes that represent these programs are shown, all significantly correlated with pseudotime. Key enrichment pathways and genes belonging to each block are indicated at right. Subcluster colors are the same as those in the pseudotime trajectory shown in Supplementary Figure S2a. **b.** Scaled, smoothed expression of select transcriptional regulators (colored lines) and canonical markers (black dashed/solid lines) across pseudotime differentiation of human tracheal secretory cells *in vivo*. The x-axis corresponds to the x-axis in **a**. **c.** Pie chart depicts proportions of all mucus secretory cells exhibiting different *MUC5AC* and *MUC5B* mucin co-expression profiles. **d.** Co-expression of common secretory markers at the mRNA level (*Left*, FISH with *SCGB1A1* in red, *MUC5B* in green) and protein level (*Right*, IF labeling with MUC5AC in red, MUC5B in green and KRT5 in white). Scale bar 25 μm. **e.** Smoking-independent correlation coefficients of *MUC5B*-correlated and *MUC5AC*-correlated genes. Genes are colored based on whether they were significantly correlated with only *MUC5B* (green), only *MUC5AC* (blue), or both (orange). Select genes are labeled. **f.** Box plots illustrate the converse effects of smoking on the mean expression of *MUC5B* and *MUC5AC*-correlated genes. P-values are from one-sided Wilcoxon tests. See also Supplementary Figure S2.

A second module followed that was characteristic of club cells^32^ including expression of *SCGB1A1*, *WFDC2*, and *CYP2F1*. This module was highly enriched for O-linked glycosylation of mucins and xenobiotic metabolism, and contained airway transmembrane mucin genes^33^ (*MUC1*, *MUC4*, *MUC16*). In this club secretory phase of pseudotime, an array of known and novel TFs increased expression, eventually reaching a crescendo in the mucus secretory cells, consistent with these cells transitioning into mucus secretory cells. The first of these to appear was the novel cAMP responsive TF, *CREB3L1*, which was followed by expression of both *SPDEF* and another novel TF, *MAGED2*. Expression of *SCGB1A1* began later and was coincident with expression of *XBP1*, a TF likely driving the cellular stress response to the initial secretion of secretoglobin and accompanying secreted proteins^34,35^ (Figure 2b). The secretory cell trajectory terminated with the mucus secretory module, containing both *MUC5AC*, *MUC5B*, and the TF, *FOXA3*, while being highly enriched for genes involved in O-linked glycosylation, vesicle coating, SLC-mediated membrane transport, and unfolded protein response, consistent with these cells actively producing and secreting mucus (Figure 2ab). Together, these data support a single developmental lineage of human secretory cells, driven by sequentially activated TFs, which transitions through functional intermediates (club cells) to culminate in a multi-functional mucus secretory cell.

### Heavy smoking drives mucus secretory cells to express a *MUC5AC* secretory program

We investigated whether transcriptionally functional subsets of mucus secretory cells exist that may carry out the known mucociliary and airway defense responsibilities of these cells. Agnostic subclustering yielded two subpopulations (Supplementary Figure S2b), one of which contained only 15% of secretory cells and was surprisingly distinguished by its expression of many known ciliated cell markers, including critical regulator *FOXJ1*^16^ (Supplementary Figure S2c). We speculated that this hybrid secretory/ciliated population was undergoing transdifferentiation, discussed later. The larger subpopulation, consisting of mucus secretory cells, exhibited no functionally distinct transcriptome-level subgroups and failed to reflect goblet cell subtypes previously reported^36^ (Supplementary Figure S2d).

To further explore the heterogeneity in this mucus secretory subpopulation, we inspected the distribution of the canonical secretory genes, *SCGB1A1*, *MUC5AC* and *MUC5B*, and observed high co-expression of all three of genes, with 94% of cells expressing *SCGB1A1* and 78% of cells expressing both *MUC5AC* and *MUC5B* (Figure 2c). Pervasive co-expression was confirmed in tracheal sections at both the mRNA and protein level (Figures 1e, 2d). These patterns are consistent with the transcriptome-wide homogeneity observed and suggest these genes all reach peak expression in this mucus secretory state.

Considering that *MUC5AC* was a core smoking response gene, we next examined whether mucin co-expression differed between nonsmokers (NS) and heavy smokers (HS). We found a sharp increase in the frequency of *MUC5AC*^+^ only cells (NS=1%, HS=10%) with heavy smoking and a corresponding decrease in both *MUC5B*^+^ only (NS=17%, HS=5%) and *MUC5AC*^−^ / *MUC5B*^−^ double negative cells (NS=8%, HS=2%; Supplementary Figure S2e), consistent with published data showing that *MUC5B* is more homeostatic and defensive^37^, whereas *MUC5AC* is more inducible and characteristic of inflammatory disease states^15,30^. Moreover, we found little overlap in genes correlated with *MUC5AC* and *MUC5B*, suggesting these mucins are associated with distinct functional programs (Figure 2e). For example, *MUC5B*-specific correlated genes encoded known secretory defense proteins, including *C3*, *CFB*, *SAA1*, *SAA2*, and *LCN2*, whereas *MUC5AC*-specific correlated genes contained a different set of defensive proteins (*MSMB*, *LYZ*, *TFF1*, *BPIFB2*, *CEACAM5*) while also being enriched for pro-secretory pathways related to ER-based protein processing and glycosylation (Figure 2e). Notably, among genes uniquely co-expressed with *MUC5AC* was *XBP1*, a TF previously implicated in both mucus production and its associated unfolded protein responses^34,38^. Not only were the two mucins themselves anti-correlated with smoking, but mean expression of *MUC5AC-* or *MUC5B*-correlated genes also increased or decreased, respectively, with smoke exposure (Figure 2f). Both *IL33*, a master regulator of type 2 mucus metaplasia^39–41^, and *NKX3-1* are potential regulators of these smoking-induced changes in secretory cells (Supplementary Figure S2f). Together, these data further support the concept of a continuous secretory cell lineage and show how smoking may mediate an additional transition, from mucin-balanced terminal secretory cells into an extended endpoint where *MUC5AC* (and its co-expressed program) dominate.

### *MUC5B*^high^ SMG secretory cells shift toward *MUC5AC* production and away from specialized defensive secretions with heavy smoke exposure

Human airway mucus is formed from the composite of secretions produced by both surface and SMG mucus secretory cells^42^. We thus compared expression profiles between these two populations to examine similarities and differences in their secretory products and the molecular mechanisms that underlie them.

We identified over 100 DEGs defining mucus secretory cells in both SMG and surface populations, which were enriched for transmembrane transport and mucosal defense (Supplementary Figure S3a). Despite these similarities, an even larger number of genes were uniquely characteristic of one or the other cell type (Figure 3a, Supplementary Figure S3b). The SMG population specifically expressed a highly unique repertoire of secretory proteins with strong enrichment for bacterial defense and innate immunity functions (Figure 3a). Furthermore, we found that while both populations highly expressed *MUC5B* (Figure 3b), expression of *MUC5AC* in SMG secretory cells was much lower (18.4-fold reduction) and less ubiquitous (SMG=30% vs. surface=84%). Reduced *MUC5AC* within SMG cells was accompanied by significantly reduced or absent expression of a host of genes involved in ER-to-Golgi vesicle-mediated transport, protein processing in the ER, and both O-linked and N-linked mucin glycosylation (Figure 3a). Distinct panels of TFs in the two groups likely govern these different expression states. For example, *CREB3L1* and *SPDEF*, the canonical secretory cell TF, were most predominant on the surface, while SMG cells uniquely expressed the SMG TF, *SOX9*, as well as *FOXC1* and *BARX2*, which are known to be involved in lacrimal gland development^43–45^ (Figure 3a). Together, our data suggest that unique TF drivers in SMG cells result in *MUC5B*-dominated mucus, which requires considerably less post-translational processing and glycosylation than surface mucus production, and is equipped with specialized defensive functions. This is consistent with recent studies detailing distinct physical properties of mucus from the SMG compared to epithelial surface^46,47^.

**Figure 3:**
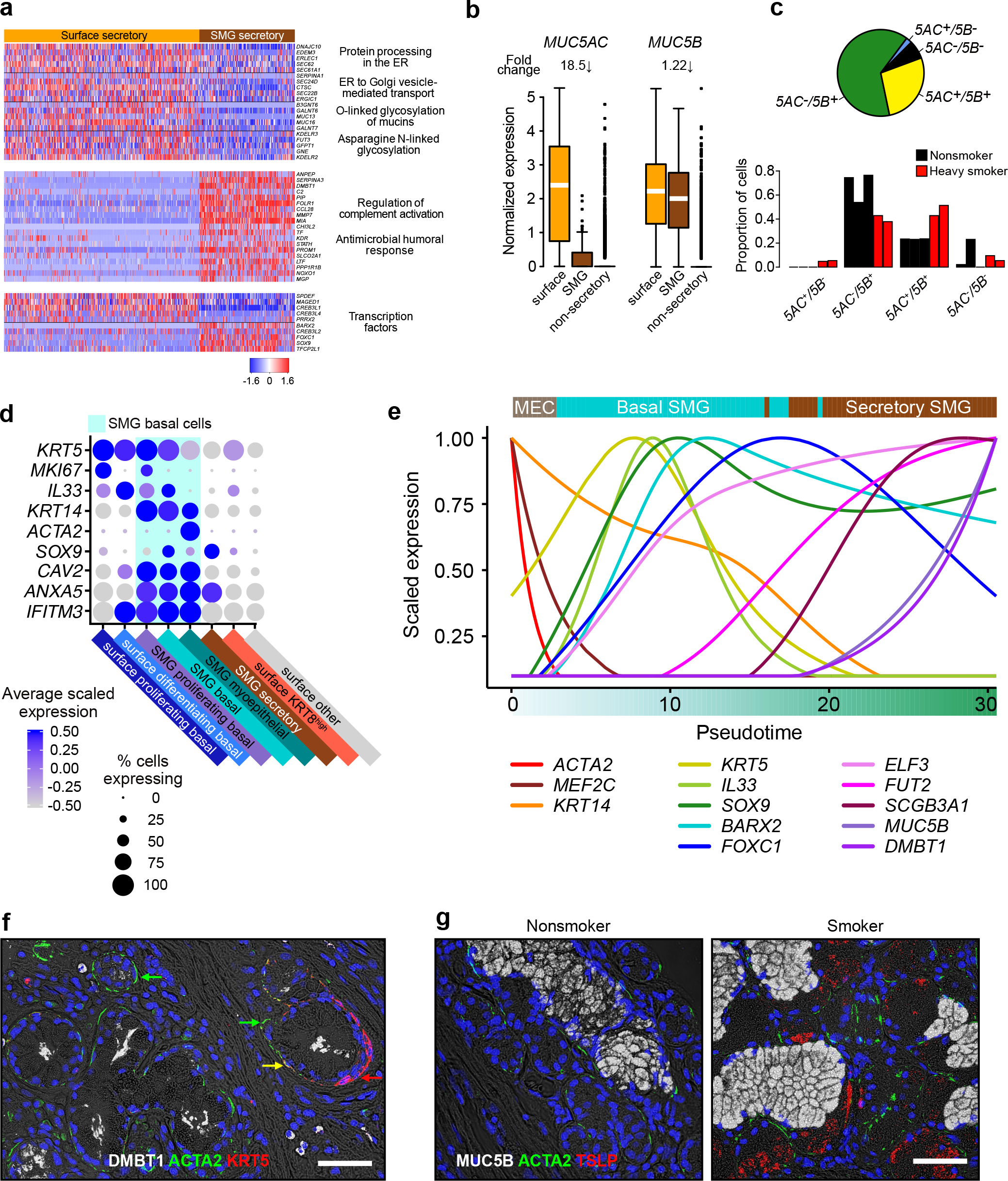
SMG mucus secretory cells are predicted to derive from a myoepithelial cell and exhibit specialized functions which are modified by smoking. **a.** Heat map depicts select genes, functional terms and TFs that distinguish surface and SMG secretory cells. Detailed heat map in Supplementary Figure S3a. **b.** Box plots of normalized mucin expression across surface secretory (orange), SMG secretory (brown) and non-secretory cells (grey). Median fold change between surface and SMG secretory cells is indicated. **c.** *Top*, Pie chart depicts proportions of SMG secretory cells exhibiting different *MUC5AC* and *MUC5B* mucin co-expression profiles. *Bottom*, bar plot showing how proportions of cells belonging to each mucin co-expression class differ between nonsmokers (black; n = 3) and heavy smokers (red; n = 2). **d.** Dot plot showing the expression of markers that unite and distinguish SMG basal cell substates, relative to surface populations and each other. **e.** Scaled, smoothed expression of key genes and regulators across a pseudotime trajectory that models the differentiation process of SMG cells. MEC, myoepithelial cells. A minimum spanning tree of the trajectory can be found in Supplementary Figure S3h. **f.** IF labeling illustrates myoepithelial cells (ACTA2^+^, green) transitioning to SMG basal cells (KRT5^+^, red). Example myoepithelial (green arrows), SMG basal (red arrow) and transitioning (yellow arrow) cells are highlighted. DMBT1 is in white and scale bar is 50 μm. **g.** IF labeling illustrates presence of TSLP (red) in SMG basal cells of heavy smokers. ACTA2, green; MUC5B, white. Scale bar, 25 μm. See also Supplementary Figure S3.

Examining smoking effects in SMG mucus secretory cells, we found that *MUC5AC* was induced and *MUC5B* was suppressed by heavy smoking (Figure 3c), echoing mucin responses on the surface. Heavy smoking also increased levels of inflammatory cytokine interleukin-6 (*IL6*) uniquely in these cells, which has been implicated in lung disease^48^ (Supplementary Figure S3c). However, most notable was the unique downregulation of 48 genes in SMG cells with smoking, which were related to multiple functions, including calcium ion binding (*S100A6*, *S100A16*) and secretion (e.g., *DMBT1*, *TF*, and *GNAS*) (Supplementary Figure S3c). Heavy smoking thus appears to induce inflammation, shift the balance of mucins, and diminish the diversity of specialized proteins produced by SMG mucus cells, likely compromising barrier and defense functions of the airway.

### Human SMG basal cell states include myoepithelial cells and are modified by heavy smoking

Recent work in mice has established the myoepithelial cell population as the SMG stem cell, which can differentiate into luminal cells through a basal cell intermediate. These cells were also shown to regenerate the surface epithelium in settings of severe injury. How these observations translate to the human airway is unclear. Investigating this, we identified a cell population with high glandular gene expression which highly expressed *KRT14*, a marker of murine SMG basal cells^2,49,50^. Also upregulated in this population were several other genes associated with glandular basal cells, including *CAV1*, *CAV2*, *IFITM3, ACTN1*, and *VIM*^51–53^ (Figure 3d, Supplementary Figure 3d).

Subclustering of this group revealed additional heterogeneity, with three major states identified (Supplementary Figure S3defg). The smallest of these expressed over 200 genes related to muscle function absent from the other two subpopulations (Supplementary Figure S3dg), a profile highly similar to murine SMG myoepithelial (*ACTA2*^+^) cells^54,55^. This population was not proliferating (*MKI67*^−^) and poorly expressed *KRT5*, suggesting it represents a quiescent human myoepithelial population. Compared to myoepithelial cells, the other two major SMG basal states exhibited expression more typical of surface basal cells, including high *KRT5* expression. One of these *KRT5*^high^ populations was proliferating (*MKI67*^+^) and in fact clustered with surface proliferating basal cells upon epithelium-wide clustering (Figure 1b). The second state was non-proliferative and appeared to be differentiating in that it highly expressed, *IL33*, a marker of surface differentiating basal cells in our dataset (Figure 3d).

A pseudotime trajectory^31^ of all (except proliferating) SMG populations proceeded from myoepithelial cells into differentiating basal, and then SMG mucus secretory cells (Figure 3e, Supplementary Figure S3h). Transitioning out of the myoepithelial state involved losing expression of *ACTA2* and muscle-related genes while simultaneously gaining expression of basal cell genes (*KRT5*, *IL33*). TFs distinctively characteristic of SMG (compared to surface) mucus cells (*SOX9*, *FOXC1*, and *BARX2*) initiated high expression in the differentiating basal population, consistent with this state being the precursor to mucus SMG cells (Figure 3e). IF labeling of tracheal sections further supported these transitions as well as the presence of these populations at the protein level (Figure 3f).

Even these SMG basal cells at the base of glands were affected by prolonged smoking, exhibiting a total of 174 DEGs (Supplementary Figure S3ij). Notably, smoker SMG basal cells uniquely upregulated *TSLP*, a major driver of type 2 airway inflammation^56,57^, which we confirmed with IF labeling (Figure 3g), suggesting a potentially unrecognized role for these cells in the onset of chronic inflammatory airway disease.

### Sequential transcriptional programs drive motile ciliogenesis

Upon cell fate acquisition, nascent ciliated cells activate expression of a large ciliary program, precipitating the generation of hundreds of cytoplasmic basal bodies which traffic to and dock with the apical membrane where they then elongate motile axonemes^58^. As our *in vivo* scRNA-seq data did not wholly capture the heterogeneity reflective of this progression, we studied the process by culturing basal tracheal epithelial cells from a subset of the donors at air-liquid interface (ALI) and harvesting replicate cultures at 20 timepoints across mucociliary differentiation for scRNA-seq sequencing analysis (Supplementary Figure S4abc). Clustering of 5,976 cells yielded three *in vitro* populations distinguished by their high expression of ciliary genes (Figure 4a, Supplementary Figure S5a). Trajectory reconstruction^59^ identified two major lineages, one of which transitioned from basal through early secretory cells, culminating in the three ciliated cell populations (Supplementary Figure S4d), whose ordering matched the real-time appearance of states across ALI differentiation (Supplementary Figure S5b). The first state to appear was highly enriched for genes involved in basal body assembly^60,61^ (*DEUP1*, *STIL*, *PLK4*) (Figure 4a) and also contained known early transcriptional drivers of ciliogenesis^62–64^ (*MCIDAS*, *MYB*, and *TP73*) (Figure 4b). We also found that TF, *E2F7*, was highly expressed in this state. Since E2F4 and E2F5 act at the top of the ciliogenesis program^62^, other family members may also be involved in this initial early ciliating stage.

**Figure 4:**
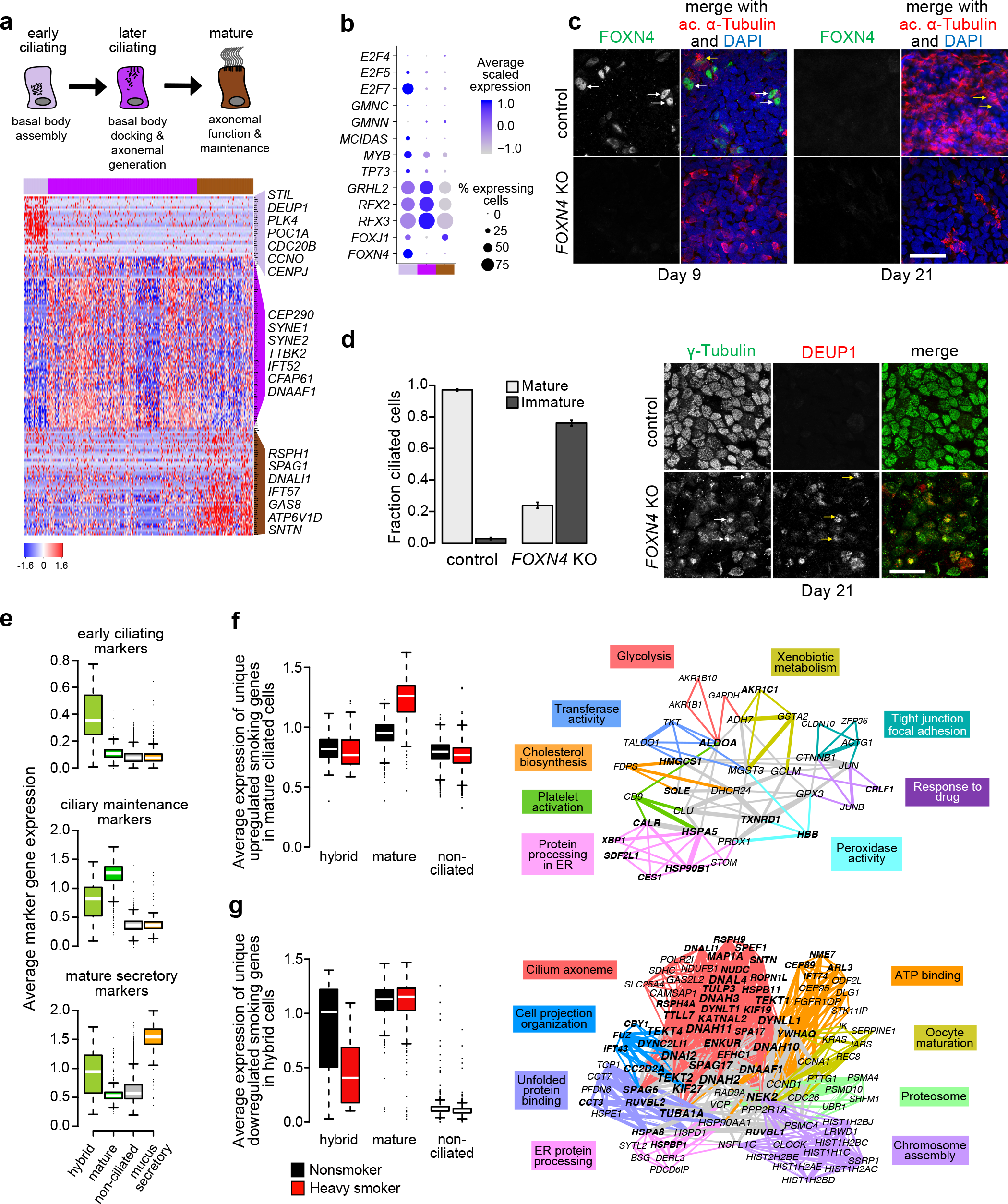
Sequential transcriptional programs drive motile ciliogenesis and smoking inhibits the early ciliating cell state. **a.** Heat map depicts gene signatures of three ciliated cell states (function summarized in schematic above) in human airway epithelial ALI cultures sampled across differentiation. Select genes from each state are indicated. **b.** Dot plots reveal TFs exhibiting expression associated with ciliated states *in vitro*. **c.** Wholemount IF labeling of *FOXN4* knockout in human tracheal epithelial ALI cultures is consistent with early expression of FOXN4 (green). Acetylated α-Tubulin (ACT), red, identifies immature (white arrows) and mature ciliated cells (yellow arrows) in control cultures at ALI timepoints indicated. Scale bar = 25 μm. **d.** *Left*, Quantification of mature and immature ciliated cells on Day 21 as determined by ACT labeling morphology (see Methods). *Right*, Wholemount IF labeling of *FOXN4* knockout illustrates aberrant ciliogenesis where basal bodies are generated (γ-Tubulin, green), but fail to dock (white arrows) and deuterosomes are assembled (DEUP1, red), but retained (yellow arrows). Scale bar = 25 μm. **e.** Average expression of markers from two *in vitro* ciliogenesis states (top and middle) and *in vivo* mature mucus secretory cells (bottom) reveals that the hybrid secretory/ciliated state contains both early ciliating and mature secretory character. Marker expression for the later ciliating *in vitro* state in Supplementary Figure S5e. **f.** *Left*, box plots summarize expression of genes uniquely upregulated in mature ciliated cells with heavy smoking, *Right*, functional gene network (FGN) of non-core upregulated genes in mature ciliated cells. Colored edges indicate shared enrichment annotations between genes that belong to functional categories summarized by the exemplar terms listed, grey edges indicate shared annotations across different functional categories. Edge thickness corresponds to the number of shared terms. Genes in bold are those uniquely upregulated in mature ciliated cells. **g.** Smoking downregulates ciliogenesis in hybrid secretory/ciliating cells but not mature ciliated cells. *Left*, box plots summarize expression of genes uniquely downregulated in hybrid ciliating cells, *Right*, FGN summarizes functional relatedness among these genes and is as described for **f** except that genes in bold are those annotated as cilia-related by CiliaCarta^82^. See Supplementary Figures S4-S5.

The two subsequent states were both highly enriched for mature ciliated cell genes, but the first of these in pseudotime was distinguished by the presence of basal body docking (*CEP290*, *TTBK2*)^65^ and axoneme assembly (*IFT52*) genes (Figure 4a), as well as peak expression of known ciliogenesis TFs (*GRHL2*, *RFX2* and *RFX3*) ^17,66,67^. These TFs were downregulated in the third and final state, which displayed the highest expression of another canonical ciliogenesis and ciliary maintenance TF, *FOXJ1*^16^ (Figure 4b). This state also showed high expression of mitochondrial formation and ATP synthesis genes, consistent with the significant energy requirements of axonemal motility^68^ (Supplementary Figure S5c). Finally, 233 known ciliary genes displayed higher unspliced-to-spliced ratios^69^ in the second compared to the third state, while only 10 genes showed the inverse pattern (Supplementary Figure S5de), supporting the trajectory’s ordering of states and illustrating a putative role of mRNA processing during the final completion of ciliogenesis.

Interestingly, forkhead box N4 (*FOXN4*), a known regulator of ciliogenesis in *Xenopus*^70^, was highly and nearly exclusively expressed in the early state, and may thus be a novel driver of this population. Consistent with its early expression, we detected nuclear FOXN4 at ALI Day 9 but no signal in mature ciliated cells at ALI Day 21 (Figure 4c). CRISPR-Cas9 knockout (KO) of *FOXN4* carried out in basal cells resulted in a partially penetrant block to ciliogenesis upon differentiation. At Day 21, 76% of ciliated cells had no or only short, sparse cilia compared to only 2% in the control (Figure 4d, left; Supplementary Figure S5f). The abnormal KO cells retained basal bodies and deuterosomes^60^ in the cytoplasm (Figure 4d, right), indicating that the basal body generation machinery was intact, but basal body docking and deuterosome disassembly was blocked. Thus, our data are consistent with *FOXN4* regulating this later step in early ciliogenesis.

*In vivo*, most ciliated cells were mature and the only cluster resembling the early ciliating state (Figure 4e, top), including high *FOXN4* expression, was the hybrid secretory/ciliated cell population that subclustered out of mature mucus secretory cells (Supplementary Figure S2cd). These hybrid cells expressed *SPDEF*, *MUC5AC*, and mature mucus secretory genes (Figure 4e, bottom), in contrast to the non-mucus producing early secretory cells that gave rise to the *FOXN4*^+^ early ciliating state *in vitro* (Supplementary Figure S4d). Together these data suggest that during *de novo* epithelization, ciliated cells derive from early secretory cells, but in the homeostatic airway, mature mucus cells transdifferentiate into ciliated cells, possibly in response to stimulus.

Mature ciliated cells exhibited 42 genes uniquely upregulated in heavy smokers which included the TF, *XBP1*, and genes involved in ER processing and unfolded protein and heat shock responses (Figure 4f). As heat shock family chaperonins were recently shown to be required for axonemal protein complex assembly^71^, prolonged smoking may enhance the ciliated cell-specific protein-folding program to counteract smoking-related protein damage and misfolding.

Smoking leads to decreased ciliary function and ciliated cell loss^72–75^, yet we found that genes downregulated by heavy smoking in mature ciliated cells were not related to ciliogenesis or ciliary function (Supplementary Table S2). However, these genes were strongly downregulated in the hybrid secretory/ciliated cells (Figure 4g), suggesting that the ciliogenesis program in this hybrid population is uniquely vulnerable to prolonged smoking. Thus, smoking may hinder the regeneration of ciliated cells rather than impairing their function once fully developed.

### Decoupled *FOXI1* and *CFTR* expression and a potential lineage relationship among rare epithelial cell types

Subclustering of the rare cell population identified three distinct groups, each expressing canonical markers of highly disease-relevant epithelial cell types: PNECs^19^ (*CALCA*), tuft cells^22,76,77^ (*POU2F3*), or ionocytes^21,36,78^(*CFTR*) (Figure 5a). We transcriptionally defined the function of these cells in humans using differential expression analyses, which confirmed highly enriched expression of Achaete-Scute family BHLH TFs, *ASCL1*, *ASCL2* or *ASCL3*, in PNECs, tuft cells or ionocytes, respectively^36,76,79^ (Figure 5b, Supplementary Figure S6a). These data confirm that ionocytes populate the human tracheal epithelium and highly express *CFTR*, as recently recognized^36,78^. On a per cell basis, ionocyte *CFTR* expression was between 17- and 467-fold higher than in other cell types, yet low frequency of these cells (average 0.2%) means only 11% of the total *CFTR* expressed by the epithelium was derived from ionocytes, whereas other more abundant cells contribute more, such as the *KRT8*^high^ population which supplied 56% of epithelial *CFTR* (Figure 5c). Exploring whether this result was due to a scRNA-seq sampling bias, we examined bulk RNA-seq data from our ALI differentiation time course, finding that *CFTR* expression began and peaked much earlier than ionocyte marker genes (Figure 5d), further supporting a significant *CFTR* contribution from other epithelial cell types.

**Figure 5:**
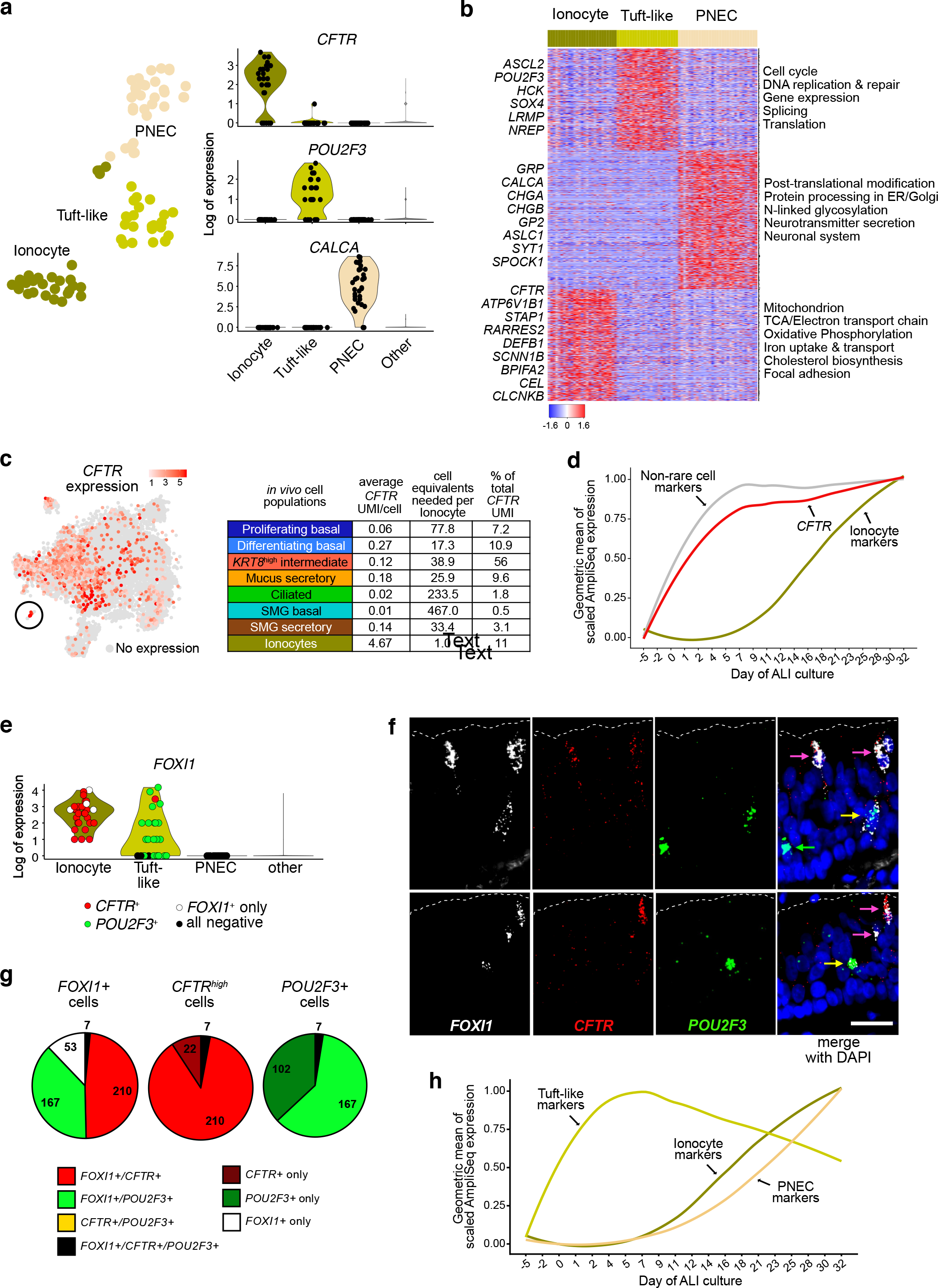
Rare cell types are highly related and *CFTR* and *FOXI1* expression is decoupled in the human airway epithelium. **a.** *Left*, tSNE depicts subclustering of rare cells found in the human tracheal epithelium, *Right*, violin plots show expression of rare cell markers identify the three subclusters. **b.** Heat map of unique gene signatures across rare cell types. Select genes in each block are indicated at left and select gene ontology terms enriched by the genes in each block are indicated at right. **c.** *Left*, Level of *CFTR* expression is shown across the tSNE plot of cells in the *in vivo* human tracheal epithelium. Grey points indicate cells with zero expression. Rare cell cluster is circled. *Right*, Table details the distribution of *CFTR* UMIs across major cell populations in the *in vivo* human tracheal epithelium. **d.** Geometric mean of scaled bulk RNA-seq expression for *in vivo* marker genes of non-rare cells, ionocytes, and *CFTR* across samples from 20 timepoints of human epithelial ALI differentiation indicate that bulk *CFTR* appears days before ionocytes in culture. **e.** Violin plots show that *FOXI1* is expressed in both tuft-like cells and ionocytes *in vivo*. Point color indicates co-expression of *FOXI1* with *CFTR* (red), *POU2F3* (green) or neither (white). **f.** FISH of *FOXI1* (white), *CFTR* (red) and *POU2F3* (green) illustrates overlap of *FOXI1* in ionocytes (*FOXI1*^+^/*CFTR*^+^, pink arrows) and tuft-like cells (*FOXI1*^+^/*POU2F3*^+^, yellow arrows) in the human tracheal epithelium *in vivo*. Green arrow, *FOXI1*^−^/POU2F3^+^tuft-like cell. Representative images from more than 8.4 cm of basolateral membrane across four donors are shown. **g.** Average co-expression quantification of FISH for the three markers in **f** across 561 total ionocytes and tuft-like cells imaged in 4 donors. Number of cells is indicated in each pie. Break down of each donor’s profile can be found in Supplementary Figure S6f. **h.** Tuft-like cell markers appear days before ionocyte or PNEC markers *in vitro*. The geometric mean of scaled bulk RNA-seq expression from the top 25 *in vivo* markers for each rare cell type are shown. See also Supplementary Figure S6.

The expression signature in our human PNECs revealed neurotransmitter processing pathway genes employed by these cells as well as a host of secreted neurotransmitters (Figure 5b). Despite characteristic *POU2F3* and *ASCL2* expression in our human tuft cell population, expression signatures in these cells were distinct from those reported in murine tuft cells^36^ (Figure 5b, Supplementary Figure S6a). For example, many diagnostic markers in mice (*GNAT3*, *TRPM5*, *GNG13*, *HMX2*, etc.) were not well-represented in our human scRNA-seq dataset, while other murine markers were present (*HOXA5*, *HCK*, and *LRMP*). Therefore, we classified our *POU2F3*^+^/*ASCL2*^+^ cells as “tuft-like” to signal the uniqueness of their transcriptional profile compared to previously described tuft cells.

Along with murine lineage tracing results^36^, the appearance of all these populations in ALI cultures (Supplementary Figure S6bcd) demonstrates that rare cells all ultimately derive from basal airway epithelial cells. Yet, little is known about the paths a basal cell takes to differentiate into these cell types. That these three rare populations clustered together when the entire epithelium was analyzed (*in vitro* and *in vivo*, and by others^78^) potentially indicates a shared origin as well as phenotype. Supporting this, differential expression analysis identified 67 genes highly expressed across only these three populations, as well as hundreds of genes uniquely shared between pairs of rare cell types (Supplementary Table S3). Interestingly, *HES6* was one of the 67 genes uniting rare cells, suggesting a possible role for NOTCH competition^80^ during fate determination of these cell types, and also potentially reflecting a shared neuronal character^81^ (Supplementary Figure S6e).

Most intriguing of the rare cell pair relationships was that observed between ionocytes and tuft-like cells, which uniquely shared expression of 114 genes, including the reported ionocyte TF, *ASCL3*^36,78^ (Supplementary Table S3). Moreover, we found that ionocyte marker, *FOXI1*^20^, whose expression has been reported to be sufficient to produce *CFTR*^high^ ionocytes^36,78^, was expressed by roughly half of *POU2F3*^+^ tuft-like cells (Figure 5e). Despite *FOXI1* expression levels comparable to ionocytes, these tuft-like cells lacked detectable *CFTR* expression. Confirming this, *FOXI1* was present by FISH in nearly all *CFTR*^high^ cells and about half of *POU2F3*^+^ cells (Figure 5f). Quantifying the FISH data, on average 48% of *FOXI1*^+^ cells exhibited an ionocyte expression pattern (*CFTR*^+^/*POU2F3*^−^), while 38% of *FOXI1*^+^ cells exhibited a tuft-like pattern (*CFTR*^−^/*POU2F3*^+^) (Figure 5g, Supplementary Figure S6f). Consistent with this, bulk RNA-seq data from the ALI differentiation time course shows that the tuft-like expression signature begins and peaks early in differentiation, whereas signatures of both ionocytes and PNECs appear much later, continuing to increase after much of the epithelium has matured (Day 21) (Figure 5h). Tuft-like cell expression was also most similar to that in basal cells (Supplementary Figure S6g), further supporting the possibility that they could serve as precursor cells. Together, these data suggest that tuft-like cells may be a precursor to ionocytes, and possibly PNECs.

We observed fewer tuft-like cells and more ionocytes in heavy smokers (Supplementary Figure S6h), suggesting that smoking may alter fate choice in favor of ionocytes over tuft-like cells and lending further evidence for a possible lineal relationship among these populations. In ionocytes, we observed 307 genes downregulated in heavy smokers, which included many genes highly specific to this cell type (Supplementary Figure S6ij), suggesting that ionocytes present in smokers, while not decreasing in number, may exhibit compromised function.

## Discussion

In this study, we have generated an agnostic atlas of the human *in vivo* tracheal airway epithelium, identifying and characterizing cell types, cell states, and lineage relationships among them. As such our study expands on the mouse *in vivo* and human *in vitro* airway epithelial atlases published recently^36,78^, and we also provide a much more densely sampled *in vitro* time course of human airway epithelial differentiation. Our data reveal that during both *in vitro* differentiation and *in vivo* homeostasis, ciliated cells derive from a secretory progenitor through multiple, discrete, transcriptional states, regulated by a suite of TFs that include *FOXN4*, which we identify as a novel regulator of the earliest ciliating state. Similarly, we show that the heterogeneity in secretory cells (club, mucus secretory cells expressing one or both of *MUC5B* and *MUC5AC*) is likely all part of a continuous secretory lineage that culminates in a multi-mucin producing mucus secretory cell.

Our atlas also produces the first transcriptional picture of human airway SMG cells, allowing us to identify a human equivalent to the recently described murine myoepithelial stem cell^54,55^. Our analysis suggests this human counterpart also exhibits stem function, as it silences its muscle expression program to assume both surface basal (*KRT5*, *TP63*) and unique glandular expression (*SOX9*), as well as engage in proliferation. This basal cell state can then differentiate into a mucus secretory cell, as orchestrated by TFs distinct from those involved in surface mucus secretory cell differentiation. The uniqueness of this program produces a vastly different secretory cell, with distinct mucin expression and processing and a specialized repertoire of protein secretions. It remains unclear whether these SMG stem cells can repopulate the surface epithelium in humans as in mice^54,55^. We confirm that the homeostatic human airway epithelium does contain ionocytes and that they highly express *CFTR*. However, the large proportion of *CFTR* expression deriving from other epithelial cell types and our observation of *FOXI1*/*CFTR* decoupling, cautions against the simple *FOXI1* -> *CFTR* -> cystic fibrosis model. Lastly, our data suggest an unrecognized lineage relationship between at least tuft cells and ionocytes, if not also PNECs, which may relate to recently reported tuft-like variants of small cell lung cancer, generally thought to be a PNEC-derived tumor^76^.

Importantly, we use scRNA-seq to deconstruct smoking effects on the epithelium to the cell type level, which we can then reassemble into a comprehensive model of how smoking modifies epithelial function as a whole (Figure 6). To summarize, pan-epithelial effects of smoking reach the basal stem cells and include induction of chemokine signaling and xenobiotic metabolism at the expense of antigen presentation and innate immune signaling. Surface and SMG secretory cells shift their mucin programs toward a *MUC5AC*-dominated inflammatory state while SMG secretory cells lose many of their distinctive defensive secretions and SMG basal cells upregulate the type 2 inflammatory cytokine, *TSLP*. Early ciliating cells preferentially lose ciliary function, potentially hindering regeneration of ciliated cells upon injury, and tuft-like cells are being depleted in conjunction with an increase in functionally-impaired ionocytes. Taken together, these data paint a smoker epithelium that has been rendered more functionally monochromatic, carrying out a *MUC5AC* inflammatory program at the expense of performing its normal defensive, interactive and reparative roles essential to lung health and homeostasis.

**Figure 6:**
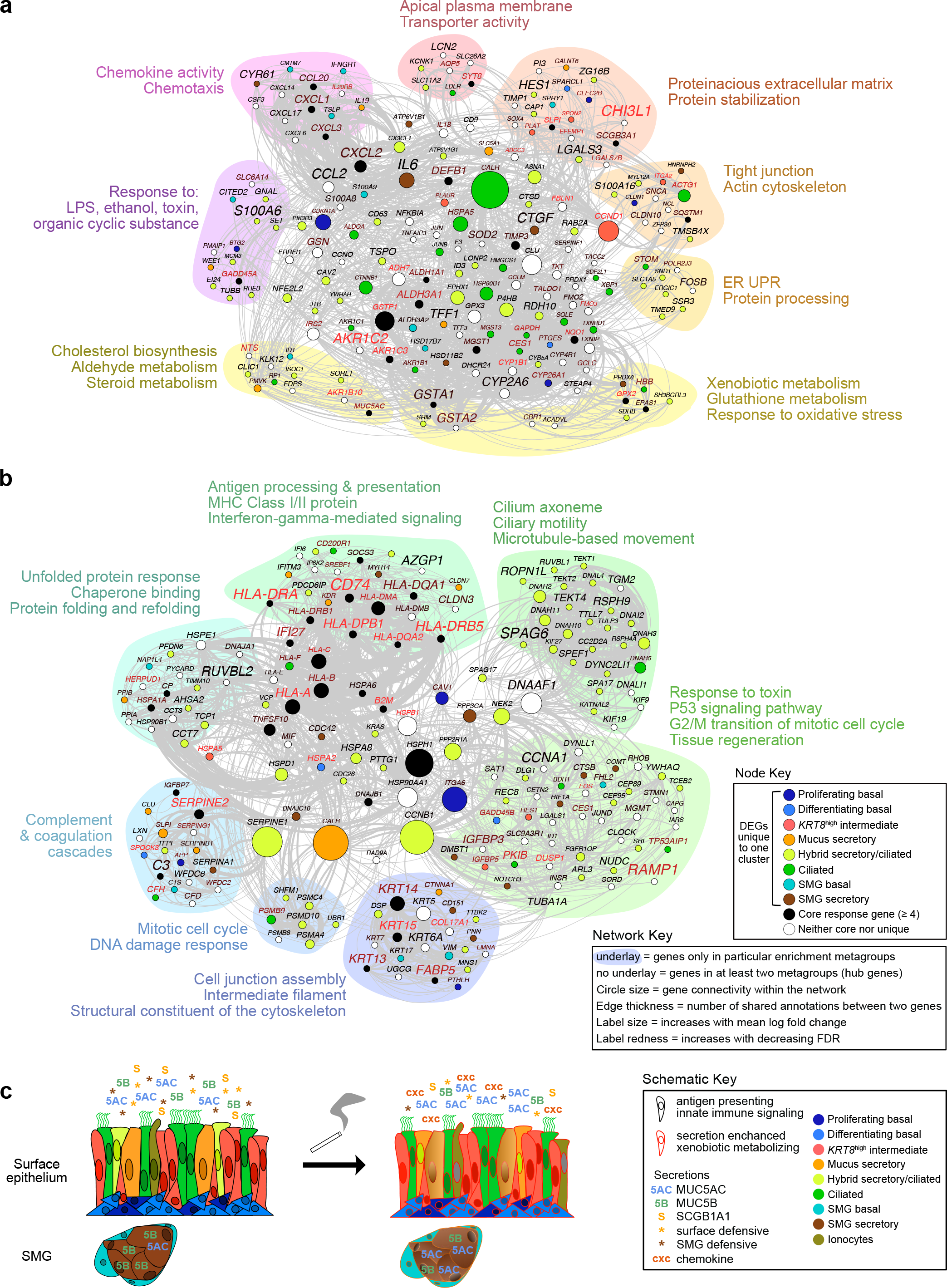
Whole epithelium smoking responses reconstructed from cell specific scRNA-seq analyses. **a.** Functional Gene Network (FGN) based on all genes upregulated with heavy smoking shows how genes that respond to smoking in distinct cell types of the airway epithelium may collaborate in carrying out dysregulated function. Node (i.e. gene) colors in Node Key refer to the cell type in which a gene was differentially expressed if “unique”; nodes for “semi-unique” and “core” DEGs are white and black, respectively. Edges connect genes annotated for the same enriched term. Exemplar enriched functions are given next to each functional metagroup (or category), which are indicated by the underlay colors that encompass all genes annotated only for the terms within the metagroup. Nodes without colored underlay represent genes in multiple metagroups. Other properties of the network, including node size, connecting edge thickness, and label size/redness are defined in the Network key. **b.** FGN as in **a**, but for all genes downregulated with heavy smoking in the airway epithelium. Legend serves for both **a** and **b.** **c.** Schematic summarizes the smoking response of the whole epithelium.

## Acknowledgements

This work was supported by the National Jewish Health Regenerative Medicine and Genome Editing Program (REGEN) and NIH grants R01 HL135156, R01 MD010443, R01 HL128439, P01HL132821, P01 HL107202. The authors would like to thank Dr. HongWei Chu, Dr. Reem Al Mubarak and Nicole Pavelka in the NJH Live Cell Core, as well as Dr. Carolyn Morris, Dr. Yingchun Li, Ari Stoner and Dr. Meghan Cromie in the Seibold Lab and Dave Heinz, Katrina Diener and Todd Woessner for assistance with tissue processing, sequencing and useful discussion.

## Author contributions

Conceptualization, K.C.G. and M.A.S.; Methodology, K.C.G., N.D.J., S.P.S. and M.A.S.; Software, N.D.J., S.P.S., N.D. and K.S.L.; Validation, K.C.G., N.D.J., E.K.V. and M.A.S.; Formal Analysis, N.D.J, S.P.S., N.D., E.G.P. and K.S.L.; Investigation, K.C.G., C.L.R., M.T.M., J.L.E. and E.K.V.; Resources, K.C.G., C.L.R., J.L.E., E.K.V. and M.A.S.; Writing – Original Draft, K.C.G., N.D.J. and M.A.S.; Writing – Review & Editing, K.C.G., N.D.J., E.K.V. and M.A.S.; Visualization, K.C.G., N.D.J., S.P.S., N.D., K.S.L., E.G.P., E.K.V. and M.A.S.; Supervision, K.C.G., N.D.J., E.K.V. and M.A.S.; Funding Acquisition, M.A.S.

## Methods

### MATERIALS AND CORRESPONDENCE

Further information and requests for resources and reagents should be directed to and will be fulfilled by Max A. Seibold

### EXPERIMENTAL METHODS

#### Key Resources

All reagents and resources referred to below are summarized with vendors and identifiers in Supplementary Table S4.

#### Human trachea samples

Human tracheal airway epithelia were isolated from de-identified donors whose lungs were not suitable for transplantation. Lung specimens were obtained from the International Institute for the Advancement of Medicine (Edison, NJ) and the Donor Alliance of Colorado. The National Jewish Health Institutional Review Board (IRB) approved the research under IRB protocols HS-3209 and HS-2240. Smokers with at least 15 pack years were classified as “heavy”, while smokers with fewer than 5 pack years were classified as “light”. See table in Supplementary Table S1 for donor details.

#### *In vivo* harvest for scRNA-seq (10X Genomics)

Human tracheas were wet in Stock solution (DMEM-F + 1X PSA) and fat and connective tissue were removed, before cutting into small sections. Sections were rinsed in Stock solution to remove mucus before proteolytic digest (0.2% Protease in Stock solution) overnight at 4°C, with rocking. Protease was neutralized with FBS, the supernatant was saved (tube 1), and tracheal sections washed (5 mM HEPES, 5 mM EDTA, 150 mM NaCl) for 20 min at 37°C. The supernatant was also saved (tube 2) and the loosened epithelium was then manually scraped off into stock solution with 10% FBS (tube 3), and all cells were collected by centrifugation for 10 min at 1000 rpm, 4°C (tubes 1, 2, and 3). Cell pellets resuspended in BEGM+0.5X PSA were filtered using a 70um cell strainer, collected by centrifugation (5 min, 1000 rpm, 4°C) and cryopreserved in freeze media (F-media, 30% FBS, 10% DMSO). On the day of capture, cells were quick thawed, washed twice in 1X PBS/BSA (0.04%) and resuspended at 1200 cells/μL for capture on the 10X Genomics platform.

#### Primary cell culture

Primary human basal airway epithelial cells from tracheal digests were expanded at 37°C on NIH 3T3 fibroblast feeders in F-media (67.5% DMEM-F, 25% Ham’s F-12, 7.5% FBS, 1.5 mM L-glutamine, 25 ng/mL hydrocortisone, 12 5ng/mL EGF, 8.6 ng/mL cholera toxin, 24 μg/mL Adenine, 0.1% insulin, 75 U/mL pen/strep) with ROCK1 Inhibitor (RI, 16 μg/mL) and antibiotics (1.25 μg/mL amphotericin B, 2 μg/mL fluconazole, 50 μg/mL gentamicin)^83^ and cryopreserved in freeze media upon initial passaging (P1).

#### *In vitro* ALI culture

Tracheal cells (P1) were expanded a second time on feeders in F-media/RI and harvested by differential trypsinization with FBS neutralization. After washing in 1X PBS, cells were resuspended in 1X HBSS and subjected to DNase digest for 5 min at 37°C. DNase was diluted 2-fold with HBSS, cells were centrifuged 1000 rpm, 5 min, 4°C and seeded onto bovine collagen-coated 6.5mm transwell inserts (2×10^4^ cells/insert) in ALI Expansion medium (50% BEBM, 50% DMEM-C, 0.5 mg/mL BSA, 80 μM ethanolamine, 10 ng/mL hEGF, 0.4 μM MgSO4, 0.3 μM MgCl2, 1 μM CaCl2, 30 ng/mL retinoic acid, 0.8X insulin*, 0.5X transferrin*, 1X hydrocortisone*, 1X epinephrine*, 1X bovine pituitary extract*, 1X gentamicin/amphotericin*, *relative to BEGM Bullet Kit aliquot) with RI (Day-5). RI was removed after 24 h, and ALI expansion medium was changed 48 hlater (Day-2). After another 48 h(Day 0), apical medium was removed and basolateral medium was replaced with PneumaCult (PC)-ALI media. Basolateral medium was exchanged for fresh PC-ALI every 48 or 72 h for the subsequent 11 days, and then daily for the following 22 days along with an apical wash of 20 μL PC-ALI.

#### *In vitro* harvest for scRNA-seq (WaferGen)

For each timepoint, medium was removed from endpoint ALI cultures, and the apical chamber was washed with warm PBS/DTT (10mM) for 5 min at 37°C, followed by a warm PBS wash of both chambers. Cultures were dislodged from the insert with 200 μL apical dissociation solution (Accutase with 5mM EDTA and 5mM EGTA) for 30 min at 37°C with occasional manual agitation^84^. Single cell suspensions were diluted, centrifuged, washed once with PBS/DTT and twice with PBS, before cryopreservation (F-media, 40% FBS, 10% DMSO). On the day of capture, cells were quick thawed, washed with 1X PBS (no BSA) and counted before proceeding with WaferGen capture according to the manufacturer’s instructions with the following modifications: 5 or 10 × 10^4^ cells were stained per sample, single cell candidates were confirmed by manual visual triage.

#### FOXN4 KO in human tracheal basal cells

Two CRISPR RNA (crRNA) guides targeting human FOXN4 annealed with universal tracrRNA and complexed with Alt-R HiFi Cas9 nuclease (1:1.2 duplex:nuclease), were electroporated (3.1uM RNP) into expanded human tracheal basal cells (5 × 10^5^ cells/transfection) with the Amaxa Nucleofector II (pulse code W-001). RNP-containing basal cells were then expanded on rat collagen-coated dishes and seeded onto transwell inserts (1 × 10^5^ cells/insert) in PC-Ex Plus expansion media with RI. RI was removed after 24 h, and 48 h later apical medium was removed and basolateral medium was replaced with PC-ALI medium. Basolateral media was replaced with fresh PC-ALI every 48 or 72 h until harvest for wholemount IF labeling on the day indicated.

#### Immunofluorescence microscopy

##### Tissue and ALI section histology

Adjacent cross-sections of human trachea were fixed in 10% neutral buffered formalin for >48 h at 4°C. ALI cultures were washed with warm 1X PBS (5 min, 37°C), fixed in PFA (1X PBS, 3.2% PFA, 3% sucrose) for 20 min on ice and washed twice with ice cold PBS. Tissue or ALI cultures were cut out of plastic supports, paraffin-embedded and sectioned onto microscope slides. Rehydration was performed with two 3 min washes in HistoChoice, followed by a standard ethanol dilution series, and antigen retrieval in Antigen Unmasking Solution for three 4-min boiling intervals. Slides were then cooled to room temperature on ice and washed three times with TBST (1X TBS, 0.5% TritonX-100) before blocking in Block buffer (1X TBS, 3% BSA, 0.1% TritonX-100) for 30 min at room temperature. Double or triple primary antibody applications were performed in Block buffer overnight at 4°C with dilutions as follows: KRT5 (1:500), TP63 (1:200), MKI67 (1:100), KRT8 (1:100), MUC5AC (1:500), MUC5B (1:200), KRT14 (1:200), DMBT1 (1:50), SCGB3A1 (1:20), ACTA2 (1:100), TSLP (1:30). Slides were washed three times in TBST before concurrent secondary application (1:500) in Block buffer with DAPI for 30 min and room temperature. Slides were again washed three times in TBST, mounted with ProLong Diamond Mount and imaged on an Echo Revolve R4 microscope.

##### ALI wholemount

ALI cultures inserts were rinsed with 1X PBS, fixed for 15 min in 3.2% PFA (no sucrose) at room temperature or 10 min in methanol at −20°C, rinsed twice more with 1X PBS and stored at 4°C. Membranes were cut out of plastic supports and placed cell-side up on parafilm in a humid chamber. After three brief washes in TBST, cells were blocked in Block buffer for 30 min at room temperature and primary antibodies were applied in Block buffer for 2 h at room temperature at the following dilutions: FOXN4 (1:50), Acetlyated α-Tubulin (1:1000), γ-Tubulin (1:500), DEUP1 (1:500). Membranes were washed 3 times in TBST before concurrent secondary application (1:500) with DAPI for 30 min at room temperature. Membranes were again washed three times in TBST, mounted with ProLong Diamond and imaged on a Leica TCS SP8 confocal microscope. Ciliated cells were quantitated based on anti-acetylated α-Tubulin antibody labeling in maximum projections of confocal image stacks of scramble (584) or FOXN4 KO (854) cells. “Mature” ciliated cells display numerous well-formed cilia, and “immature” cells display one of the indicated phenotypes: short, sparse or bulging.

#### Fluorescence *in situ* hybridization (RNAScope)

Adjacent cross-sections of human trachea or PBS rinsed ALI cultures were fixed in 10% neutral buffered formalin for 24+/− 8 h at room temperature, washed with 1X PBS and paraffin-embedded immediately. Paraffin blocks were sectioned onto SuperFrost Plus slides, dried overnight and baked for 1 h at 60°C before immediately proceeding with the RNAScope Multiplex Fluorescent v2 assay according to the manufacturer’s instructions with the following modifications: Target retrieval was performed for 15 min in a boiling beaker, protease III was used for 30 min pre-treatment at 40°C and hybridized slides were left overnight in 5X SSC before proceeding with the amplification steps. Opal fluors were applied at 1 in 1500 dilution, and slides were image on an Echo Revolve R4 microscope. Ionocytes and tuft-like cells were quantified by presence of grouped *FOXI1*, *POU2F3* and/or *CFTR* puncta on triple labeled sections. Basolateral membrane length was quantified with the freehand line tool and Measure functions in ImageJ.

#### AmpliSeq of bulk RNA samples

Bulk RNA was extracted with the Quick RNA MicroPrep Kit and 0.5 – 3 × 10^5^ cells were harvested alongside those used for scRNA-seq. Isolated RNA was normalized to 3.5 ng input/sample for automated library preparation with the Ion AmpliSeq Transcriptome Human Gene Expression Kit, using 12 cycles of amplification. Libraries were sequenced with Ion Proton System (ThermoFisher Scientific).

### QUANTIFICATION AND STATISTICAL ANALYSIS

#### Pre-processing of *in vivo* scRNA-seq data

Initial pre-processing of the 10X *in vivo* scRNA-seq data, including demultiplexing, alignment to the hg38 human genome, and UMI-based gene expression quantification, was performed using Cell Ranger (version 2.1, 10X Genomics).

We next carried out donor-specific filtering of cells to ensure that high quality single cells were used for downstream analysis. Although samples comprising multiple cells were removed during the cell selection stage, we safeguarded against doublets by removing 465 cells with either a gene count over the 99th percentile or a UMI count over the 97th percentile. Furthermore, 223 cells were removed that either exhibited fewer than 1500 genes or fell within the 1st percentile of gene counts and 170 cells were removed that contained more than 30% of mapped reads originating from the mitochondrial genome. Prior to downstream analysis, select mitochondrial and ribosomal genes (genes beginning with MTAT, MT-, MTCO, MTCY, MTERF, MTND, MTRF, MTRN, MRPL, MRPS, RPL, or RPS), or very lowly expressed genes (expressed in < 0.1% of cells) were also removed. The final quality-controlled dataset consisted of 14,324 cells, and 19,339 genes. After initial clustering and visualization, which allowed for identification of 11 major cell populations (see Supplementary Figure S1a), 484 cells were removed that we characterized as non-epithelial (smooth muscle cells, endothelial cells, and immune cells), leaving 13,840 epithelial cells for further analysis.

To account for differences in coverage across cells, UMI counts were normalized for each donor, dividing each count by the sum of its cell’s UMIs, multiplying by 10,000, and then taking the natural log. For input into dimensionality reduction and clustering analyses and for plotting relative expression in heat maps and dot plots, we also fitted normalized expression of each gene to the sum of the UMI counts per cell, and then mean centered (subtracted from each gene count the average expression of that gene) and scaled (divided each centered gene count by that gene’s standard deviation) the residuals. The Seurat R package^85^ was used to carry out all data normalization and scaling as well as downstream dimensionality reduction, clustering, tSNE plot overlaying, and differential expression.

#### Pre-processing of *in vitro* scRNA-seq data

We trimmed and culled raw demultiplexed cDNA reads in FASTQ files using Cutadapt^86^, trimming poly A tails and 5’ and 3’ ends with q < 20 and removing any reads shorter than 25 base pairs. Trimmed reads were then aligned to the hg38 human genome with GSNAP^87^, setting “max-mismatches=0.05” and accounting for both known Ensembl splice sites and SNPs. Gene expression was quantified using HTSeq^88^ with “stranded=yes”, “mode=intersection-nonempty”, and “t=gene” and then summed the number of unique molecular identifiers (UMIs) for each gene across runs for each cell to obtain a UMI count matrix used for all downstream analysis.

We carried out quality-control filtering using a similar approach to that used for the *in vivo* dataset, removing 100 cells for which the percentage of reads mapping to genes was <50%, 2,262 cells with > 25% of mapped reads being mitochondrial, and 384 cells with UMI counts outside the 3rd and 97th percentiles. We filtered genes using the procedure described above for the *in vivo* dataset. The final quality-controlled dataset consisted of 5,976 cells, and 23,825 genes.

#### Pre-processing of *in vitro* bulk AmpliSeq data

Reads sequenced on the Ion Torrent Proton sequencer were mapped to AmpliSeq transcriptome target regions with the torrent mapping alignment program (TMAP) and gene count tables were generated for uniquely mapped reads using the Ion Torrent ampliSeqRNA plugin. After removing duplicated sequences our dataset contained 20,869 genes for 20 sampled time points. We normalized expression based on size factors calculated using DESeq2^89^.

#### Dimensionality reduction, clustering, and visualization

Prior to clustering and visualizing each of the two scRNA-seq datasets (*in vitro* and *in vivo*), we reduced the dimensionality of variation in a way that accounted for batch-based shifts in expression among donors. To do this, we used canonical correlation analysis (CCA) to identify the strongest components of gene correlation structure that were shared across donors (using Seurat’s RunMultiCCA function). For the *in vivo* dataset, this was based on the 5,009 genes contained within the union of the top 8,000 most informative genes from each donor that involved at least five of them, where “informativeness” was defined by gene dispersion (i.e., the log of the ratio of expression variance to its mean) across cells, calculated after accounting for its relationship with mean expression (using the FindVariableGenes function). Correlated expression across donors based on the top 25 CCA dimensions was then projected into a common subspace using Seurat’s AlignSubspace function, allowing us to identify and visualize universal populations of cells, rather than those driven by technical batch or donor-specific variation. We further reduced dimensionality of these 25 subspace-aligned CCA dimensions using the Barnes-Hut implementation of t-distributed neighborhood embedding (tSNE) and then plotted cell coordinates based on the first two dimensions (perplexity = 100). We further carried out unsupervised clustering by first constructing a shared nearest neighbor (SNN) graph based on k-nearest neighbors (k = 30) calculated from the aligned CCA dimensions. The number and composition of clusters were then determined using a modularity function optimizer based on the Louvain algorithm (resolution = 0.3).

A similar approach was followed for the *in vitro* dataset, except that the union of the top 10,000 most informative genes involving two or more of the three donors was used for CCA (9,542 genes total), while SNN clustering (resolution = 0.4, k = 15) based on an SLM optimizer and tSNE visualization (perplexity = 80) were performed using the top 20 subspace-aligned CCA dimensions. Specifications for additional subclustering and visualization performed in the study are summarized in Supplementary Table S5.

#### Plotting expression across cells

For overlaying expression onto the tSNE plot or dot plots for single genes or for the average across a panel of genes, we plotted normalized expression along a continuous color scale. All heat maps showing gene expression across cells (except Figure 2a and Supplementary Figure S3h, which were created using Monocle) were produced using Heatmap3^90^ and also show scaled normalized expression along a continuous color scale, with break scales set as indicated.

#### Differential expression analysis

Differential expression for each gene between various groups specified in the text was tested using a non-parametric Wilcoxon rank sum test carried out with Seurat. We limited each comparison to genes exhibiting both an estimated log fold change > +0.25 and detectable expression in > 10% of cells in one of the two clusters being compared. We corrected for multiple hypothesis testing by calculating FDR-adjusted p-values.

Genes were considered to be differentially expressed when FDR < 0.05.

#### Functional enrichment analysis

We tested for gene overrepresentation of all target lists within a panel of annotated gene databases (Gene Ontology [GO] Biological Process [BP] 2017, GO Molecular Function [MF] 2017, GO Cellular Component [CC] 2017, Kyoto Encyclopedia of Genes and Genomes [KEGG] 2016, and Reactome 2016) using hypergeometric tests implemented with Enrichr^91^, as automated using the python script, EnrichrAPI (https://github.com/russell-stewart/enrichrAPI). We report only terms and pathways that were enriched with FDR < 0.05.

#### Identification and plotting of cell type-specific markers

To identify cell type-specific markers for both the *in vitro* and *in vivo* datasets, we first carried out pairwise differential expression analysis between each of the major clusters, downsampling large clusters to the median cluster size. Markers for each cluster were those genes exhibiting significant upregulation (FDR < 0.05) when compared against all other clusters. Markers for each cluster were sorted by FDR as calculated based on largest p-value observed for each gene across comparisons.

For plotting expression of *in vivo* rare cell types across differentiating AmpliSeq bulk samples *in vitro*, we first obtained *in vivo* cell type markers for each of the three rare cells by isolating genes that were both significantly upregulated in each rare cell type relative to one another (with FDR < 1e-5) and when compared to all non-rare cells (with FDR < 1e-5). We then plotted the geometric mean of log-normalized expression in bulk across the top 25 *in vivo* markers for each cell type (based on FDRs in the non-rare cell comparisons), after scaling values to be between zero and one (see Figure 5h). We added a small value (0.01) to expression to avoid calculating the geometric mean with zeros. For the expression of non-rare *in vivo* cluster markers shown in Figure 5d, we took the geometric mean of log-normalized expression across the top 25 markers for each of the main non-rare *in vivo* cell clusters.

For the marker tSNE overlay plots in Supplementary Figure S4e, for each *in vitro* and *in vivo* cluster, we calculated average expression across the top 100 markers (or as many as were available, if fewer than 100) and then used these values to show characteristic expression of select *in vitro* cell types on the *in vivo* cells and vice versa. Because the Secretory cell 1 *in vitro* cluster was transitionary and consequently had only a single marker, we used the top 25 most distinct genes for this population, despite these markers not being significant by FDR < 0.05. Cells on the tSNE plots were identified as being characteristic of a given cell type if marker mean expression for that cell type was at least in the 85^th^ percentile while marker mean expression for all other cell types was below the 95^th^ percentile. To render the *in vitro* Secretory cell 1 and 2 populations more distinguishable, we reduced stringency for expression of Secretory cell 1 markers to the 75^th^ percentile and increased stringency for expression of Secretory cell 2 markers to the 95^th^ percentile.

#### Defining core and unique smoking response genes

Core smoking response genes were defined as those significantly differentially expressed in heavy smokers compared to nonsmokers in four or more of the main *in vivo* populations. For this analysis, we excluded the rare cell population, which generally contained too few cells to detect significant smoking DEGs, and used a mucus secretory population from which the small subpopulation of hybrid secretory/ciliated cells was removed, as the response in this small population was so distinct. Unique response genes were those that were significantly differentially expressed in only one population while exhibiting either a log fold change < 0.25 and/or an FDR > 0.2 in all other populations. Populations assessed for uniqueness were the two basal populations, *KRT8*^high^, ciliated, SMG basal and SMG secretory populations, the hybrid secretory/ciliated subpopulation, surface secretory cells (with the hybrid subpopulation removed), and ionocytes (there were too few of the other two rare cell types to assess smoking response). Genes responding to heavy smoking in a least one cell population but not considered unique or core were defined as semi-unique.

#### Calculating correlations with *MUC5AC* and *MUC5B*

To find genes in surface secretory cells (excluding the hybrid secretory/ciliated subpopulation) whose expression was correlated with that of *MUC5AC* or *MUC5B*, we used Spearman partial correlation analysis, which calculated gene correlations while controlling for differences in expression due to smoking. Light smoker cells and cells that did not express both *MUC5AC* and *MUC5B* were excluded from this analysis.

#### Lineage trajectories

For the *in vivo* dataset, we constructed lineage trajectories for *KRT8*^high^ and mucus secretory cell populations combined and for SMG cells (myoepithelial, SMG differentiating basal, and SMG secretory populations) to better understand the genes and processes that regulate and transition across these two lineages. For the *KRT8*^high^-to-secretory cell trajectory, we first aligned donors from these two populations using the strategy outlined above and re-clustered them (alignment and clustering specifications are given in Supplementary Table S5), resulting in nine clusters. One of these clusters corresponded to a hybrid secretory/ciliated population, which we removed prior to trajectory construction. Then, using Monocle v2.8^31^, we carried out dimensionality reduction using the DDRTree algorithm, regressing out both donor identity and the number of genes per cell, and then ordered cells along a trajectory of pseudotime (using Monocle’s orderCells function; see Supplementary Figure S2a) based on their expression across the 3,000 most differentially expressed genes (sorted by q-value), as inferred using tests of gene differential expression as a function of cluster membership (using Monocle’s differentialGeneTest function). We then tested each gene for differential expression as a function of pseudotime, hierarchically clustered the significantly correlated genes (with q-value < 0.05), and then used the plot_pseudotime_heatmap function to plot smoothed scaled expression of genes belonging to four major modules across cells sorted by pseudotime, assuming that the most basal-like cells occupy the initial state (see Figure 2a). To view the expression of key regulators across the trajectory on a shared scale, we normalized all smoothed expression values to be between zero and one, and then plotted these normalized expression curves across pseudotime (see Figure 2b).

The same approach was followed for constructing the SMG cell trajectory. To order the subclustered cells (see Supplementary Figure S3d), we used the top 3,000 genes that were most differentially expressed as a function of cluster membership to one of the three SMG basal substates, the myoepithelial state, or the SMG secretory population (proliferating basal SMG cells were excluded from this analysis). Smoothed expression for 14 (hand-sorted) modules of genes that were significantly associated with pseudotime were plotted across the trajectory, assuming that the myoepithelial cells occupy the root state (see Supplementary Figure S3h).

Additionally, we constructed lineage trajectories for the *in vitro* dataset, allowing us to capitalize on the known real-time appearance of cell states across differentiation of ALI cultures. Applying the previously calculated tSNE dimensions, we used Slingshot^59^ to build lineages of cells that link *in vitro* SNN cell clusters by fitting a minimum spanning tree (MST) onto the clusters. When constructing these lineages, we only used differentiating basal, secretory and ciliating/ciliated populations, where lineages were constrained to begin with the differentiating basal population and to end with either the mature secretory (Secretory cell 2) or mature ciliated populations. We inferred two major lineages, one defining the transition from differentiating basal to Secretory cell 1 then Secretory cell 2, and the other defining the transition from differentiating basal to early and late ciliating cells, and then on to mature ciliated cells. Pseudotime values for cells were obtained for each lineage by projecting cells onto smoothed lineages constructed using Slingshot’s simultaneous principle curves method. For the two lineages, we then plotted smoothed scaled expression (as a weighted average across a 100 cell window) of select genes that were significantly associated with pseudotime based on Monocle’s differential gene test (q-value < 0.05) (see Supplementary Figure S4d).

#### Spliced/unspliced ratios

It has been shown that unspliced and spliced mRNA molecules capture earlier and later expression states, respectively, thus providing temporal information that is distinct from expression of combined RNA-seq data^92^. Thus, to further test the polarity of the later ciliating and mature ciliated expression states in the ALI cultured scRNA-seq dataset, we used the Velocyto pipeline^92^ (applying default options) to identify unspliced and spliced mRNA reads for each gene in the original *in vitro* BAM files.

#### Gene networks

We constructed functional gene networks (FGNs) in order to summarize the major processes being carried out by selected gene sets in a way that shows the genes involved and their interconnectivity. FGNs were created by finding enriched terms for the given gene set (based on Gene Ontology and KEGG Pathway libraries), filtering and consolidating these enrichments into categories (i.e. metagroups) using GeneTerm Linker^93^, and then constructing gene networks based on select metagroups using FGNet^94^, which connects genes via edges with shared annotations that fall within a particular metagroup. Genes (i.e., nodes) uniquely involved in distinct processes (i.e., metagroups; each with a different colored border) can be distinguished from those involved in multiple processes (nodes belonging to multiple metagroups, indicated with white borders). Edges indicate at least one shared annotation.

Protein-protein interaction (PPI) networks were also constructed for some gene sets, enabling us to visualize how genes within sets may produce proteins that interact within the cell to carry out particular functions. We used STRNGdb^95^ to create PPI networks, where edges are drawn between gene nodes that have predicted interactions, and where edge thickness is proportional to the predicted interaction strength. Networks were split into densely connected sub-networks containing three or more genes using the cluster_leading_eigen or cluster_edge_betweenness functions in iGraph^96^.

### DATA AND SOFTWARE AVAILABILITY

Gene lists associated with Figure and Supplementary Figure panels can be found in Supplementary Table S6. All raw and processed scRNA-seq data used in this study are in the process of being deposited in the National Center for Biotechnology Information/Gene Expression Omnibus (GEO).

