## Supplementary Information for "Dissecting the cellular specificity of smoking effects and reconstructing lineages in the human airway epithelium"

*Authors/Affiliations*

Katherine C. Goldfarbmuren[1#], Nathan D. Jackson[1#], Satria P. Sajuthi[1], Nathan Dyjack[1], Katie S. Li[1], Cydney L. Rios[1], Elizabeth G. Plender[1], Michael T. Montgomery[1], Jamie L. Everman[1], Eszter K. Vladar[3,4], Max A. Seibold[1,2,3,*]

[1]Center for Genes, Environment, and Health, National Jewish Health, Denver, CO, 80206 USA; [2]Department of Pediatrics, National Jewish Health, Denver, CO, 80206 USA; [3]Division of Pulmonary Sciences and Critical Care Medicine and [4]Department of Cell and Developmental Biology, University of Colorado-AMC, Aurora, CO, 80045 USA, [#]these authors contributed equally to this work, *correspondence:

### Supplementary Figure S1

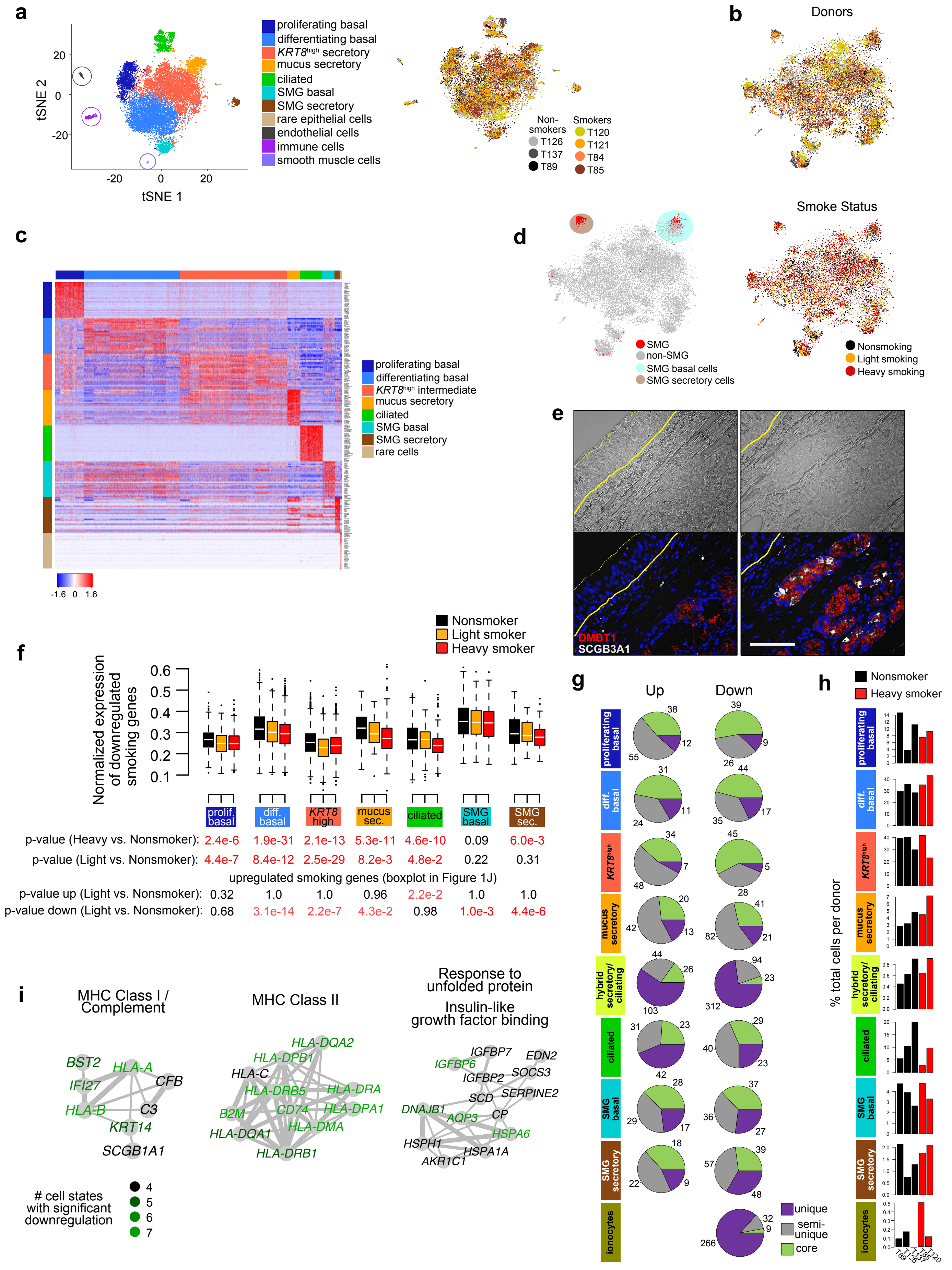

**Supplementary Figure S1 (with Figure 1): Full *in vivo* scRNA-seq analysis**

- **a.** tSNE of full dataset including three rare, non-epithelial clusters (circled). *Left,* Coloring corresponds to the cell categories indicated. *Right,* Coloring corresponds to donor/smoke status.
- **b.** tSNE of epithelial cell clusters after removing non-epithelial clusters. *Top,* Coloring corresponds to donor as indicated in legend in **a**. *Bottom,* Coloring corresponds to three-tier smoke status
- **c.** Heat map depicts gene signatures for broad cell categories in the human trachea. x-axis shows cells, y-axis gives scaled expression for the top 30 markers from each broad cell category (see Figure 1).
- **d.** The top 50 genes most upregulated in SMG compared to surface epithelial cells in Fischer et al. (2007)[1] are most expressed in our basal and secretory SMG populations *in vivo*. Red depicts cells with average expression of these 50 genes > $97.5^{th}$ percentile. Clusters classified as SMG secretory and SMG basal cells are indicated by brown and teal, respectively.
- **e.** Additional IF labeling of SMG markers, DMBT1 and SCGB3A1. Upper panels, transillumination; solid yellow line, basement membrane; dashed yellow line, luminal surface of the epithelium. Scale bar is 100 µm.
- **f.** Published bulk RNA-seq smoking downregulated genes[2] were downregulated in most cell types from heavy smokers. Box plots depict mean normalized expression for the published downregulated genes as a function of smoking habit. The first two rows of p-values are from one-sided t-tests comparing mean expression for heavy vs nonsmokers or light vs nonsmokers, as indicated for each cluster. The bottom two rows of p-values are from one-sided t-tests comparing mean expression of the published upregulated smoking genes (see Figure 1i) between light smokers and nonsmokers, where the upper values are from tests for greater means in light smokers and lower values are from tests of decreased means in light smokers. Rare cells were excluded from this analysis due to small cell numbers. Cluster colors match legend descriptions in **c**.
- **g.** Pie charts showing the proportions genes significantly up or downregulated with heavy smoking for each cluster that were core, unique, or semi-unique. Raw numbers of genes in each category are given for each category.
- **h.** Bar plots depict % of total cells for each donor belonging to each of the indicated cell clusters**.**
- **i.** PPI networks among core genes downregulated with heavy smoking.

#### Supplementary Figure S2

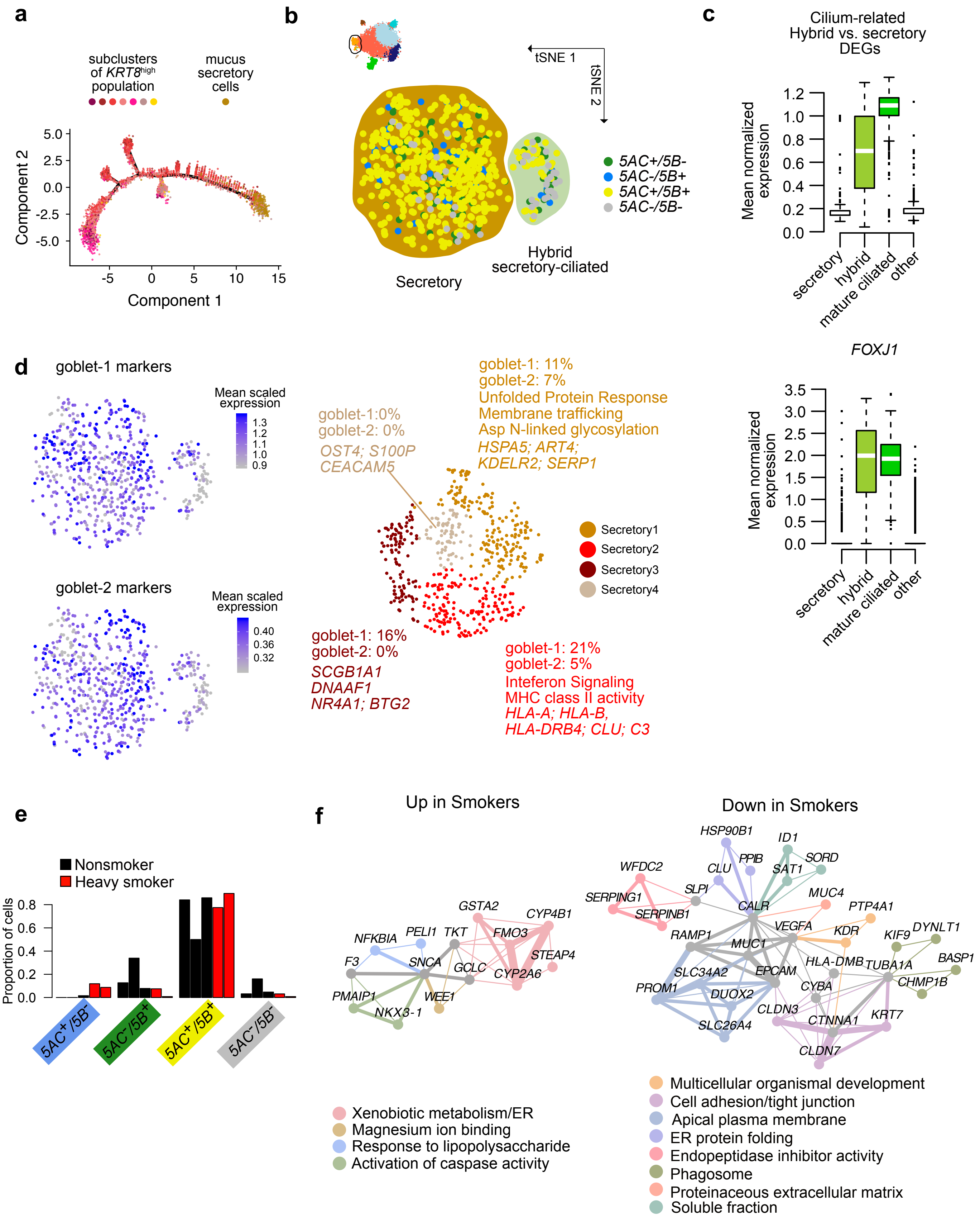

**Supplementary Figure S2 (with Figure 2):** ***In vivo*** **secretory cell lineages and smoking effects**

- **a.** Monocle pseudotime trajectory[3] of *KRT8*$^{high}$ and mucus secretory cells. Colors correspond to distinct SNN subclusters inferred for the combined dataset.
- **b.** Agnostic subclustering of the *in vivo* human secretory cells reveals a hybrid secretory/ciliated cell population (see Supplementary Table S5 for clustering specifications). Overlaid are the co-expression status of *MUC5AC* and *MUC5B*.
- **c.** Average expression of hybrid secretory/ciliated signature genes and *FOXJ1* expression illustrates the ciliated character of the hybrid population. *Top,* DEGs up in the hybrid population compared to mature secretory cells are also highly expressed in mature ciliated cells, *Bottom, FOXJ1* expression is comparable between the hybrid population and mature ciliated cells.
- **d.** Goblet cell 1 ("goblet-1") and goblet cell 2 ("goblet-2") subtype genes reported in Montoro et al. (2018)[4] do not discriminate among *in vivo* human tracheal secretory cells. *Left,* Average expression of goblet-1 (top left) and goblet-2 (bottom left) markers overlaid on a tSNE plot of only mature secretory cells from the *in vivo* human tracheal epithelium (same tSNE plot as in **c** below, see Supplementary Table S5 for specifications on how this tSNE plot was created). *Right,* Subclustering of mature secretory cells (excluding the hybrid secretory/ciliated subpopulation) using the goblet-1 and goblet-2 genes. We aligned donors using the top two CCA dimensions based on these genes and then carried out tSNE visualization and SNN clustering (SLM algorithm, resolution = 0.3, perplexity = 80). Colors correspond to four inferred clusters. The proportions of goblet-1 and goblet-2 marker genes that were among the DEGs for each of the four clusters (compared to the other three) are indicated, along with select DEGs and gene ontology terms enriched among DEGs that distinguish these clusters.
- **e.** Bar plots show differences in *MUC5AC* and *MUC5B* co-expression proportions as a function of smoking status, suggesting that double negative cells tend to acquire *MUC5AC* with prolonged exposure to smoke.
- **f.** Functional gene networks (FGN) depict interactions among genes uniquely or semi-uniquely up (left) or downregulated (right) in mature secretory cells (excluding the hybrid population) due to heavy smoking. FGNs were produced by requiring at least three genes for each enrichment term or metagroup.

#### Supplementary Figure S3

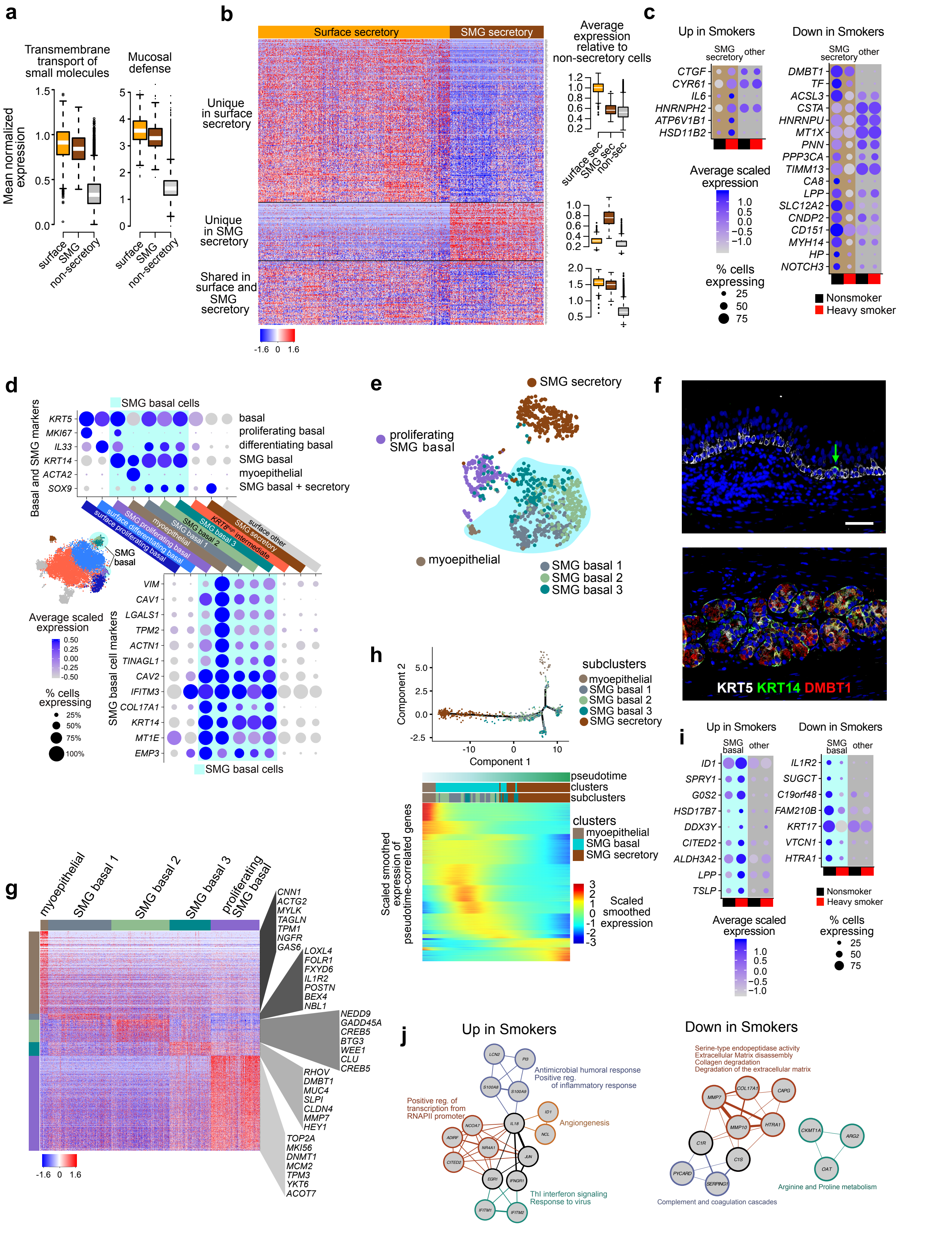

**Supplementary Figure S3 (with Figure 3): Human submucosal gland distinctions from the surface epithelium, lineages, and smoking effects**

- **a.** Box plots of selected shared functional terms between surface epithelial and SMG secretory cells. "Mucosal defense" is the average expression of the genes *BPIFB1*, *BPIFA1*, *PIGR*, *SLPI*, *LYZ*, and *WFDC2*.
- **b.** Heat map depicts all shared and unique genes significantly upregulated in surface and SMG secretory cells *in vivo* compared to non-secretory cells. Box plots at right illustrate the specificity of the heat map blocks relative to the remaining non-secretory tracheal epithelium.
- **c.** Dot plots illustrate significantly up and downregulated DEGs specific to smokers' SMG secretory cells
- **d.** *Top,* Dot plot of common basal and SMG markers across all SMG basal cell substates (highlighted in teal). *Bottom,* Dot plot of shared DEGs significantly up in all SMG basal cell states.
- **e.** tSNE of subclustered SMG cells, teal underlay specifies cells belonging to the SMG basal cell cluster shown in Figure 1b, while the proliferating SMG basal cells were originally classified as a subcluster of (surface) proliferating basal cells (see inlay in **d**). See Supplementary Table S5 for clustering specifications.
- **f.** Immunohistochemistry of SMG secretory and basal markers indicates that DMBT1 and KRT14 are highly specific to the SMG secretory and basal cells, respectively. Green arrow in top panel indicates a lone KRT14 cell in the surface epithelium.
- **g.** Heat map of DEGs distinguishing SMG basal cell states. Select genes are indicated for each block and full gene lists are present beside each row with zoom in.
- **h.** *Top,* Monocle pseudotime trajectory[3] connecting myoepithelial cells to SMG secretory cells via SMG basal cells. *Bottom,* Heat map of all genes significantly correlated with pseudotime (where myoepithelial cells were the specified root).
- **i.** Dot plots indicate up and downregulated smoking response in SMG basal cells
- **j.** Networks depict enriched functional terms connecting up- (left) and down-regulated (right) heavy smoking DEGs observed in SMG basal cells. Both unique and semi-unique DEGs were considered. Networks were created by drawing edges between SMG smoking response genes that shared one or more of the specified enriched terms, with edge thickness proportional to the number of terms shared. Terms were assigned to groups based on functional similarity which are indicated by circle border color, with black borders indicating genes annotated for enrichments in two or more groups.

#### Supplementary Figure S4

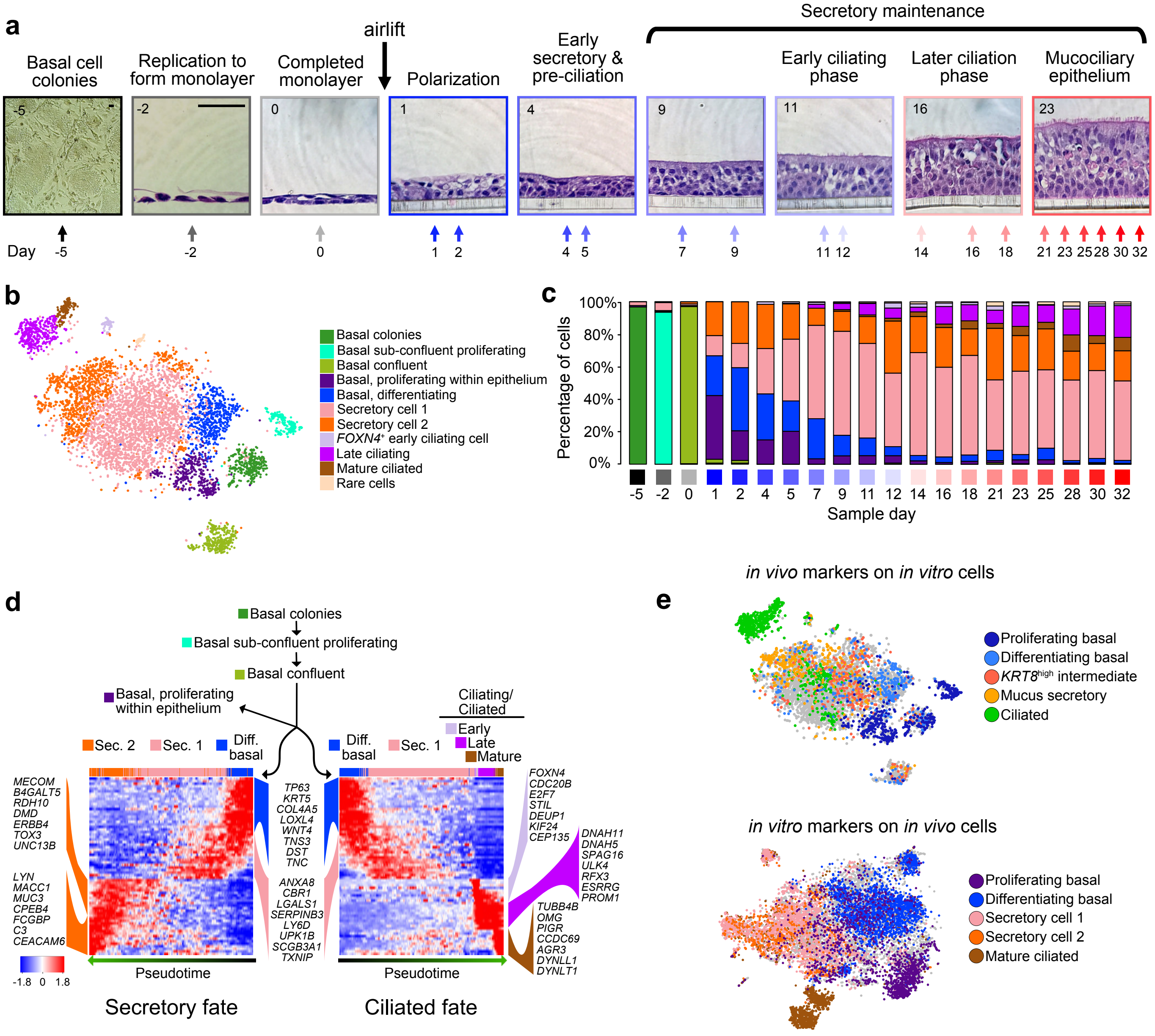

**Supplementary Figure S4 (with Figure 4): Human ALI culture model informs transitions between cell states seen in the *in vivo* human trachea**

- **a.** Histological overview of human basal cell ALI mucociliary differentiation. Image outlines: shades of black = submerged culture, shades of blue to red = polarized differentiation and maintenance of ALI human epithelial cell culture. Representative brightfield or H&E stained images of indicated time points are shown, scale bars in far left panels are both 50 μm.
- **b.** tSNE plot depicts the distribution of inferred clusters of *in vitro* cells transcriptionally sampled from across the entire differentiation time course. Cluster identities based on expressed markers are shown at the right.
- **c.** Proportion of cells in each cell state (corresponding to clusters in **b**) present at each time point over differentiation. Time course black/blue/red gradient coloring at bottom corresponds to colors in **a**.
- **d.** Similarities and differences in transcriptional programs between distinct pseudotime lineages constructed with Slingshot[5] that lead to mature secretory and ciliated cells *in vitro*. Select markers or genes correlated with pseudotime are indicated.
- **e.** Broad cell populations *in vivo* are analogous to broad cell populations *in vitro*. *Top,* characteristic expression of *in vivo* markers for each select broad cell type is overlaid onto *in vitro* tSNE cells. *Bottom,* characteristic expression of *in vitro* markers of select cell types is overlaid onto *in vivo* tSNE cells. Grey cells indicate those not characteristic of any of the broad cell types shown, based on the thresholds of expression used (see Methods).

#### Supplementary Figure S5

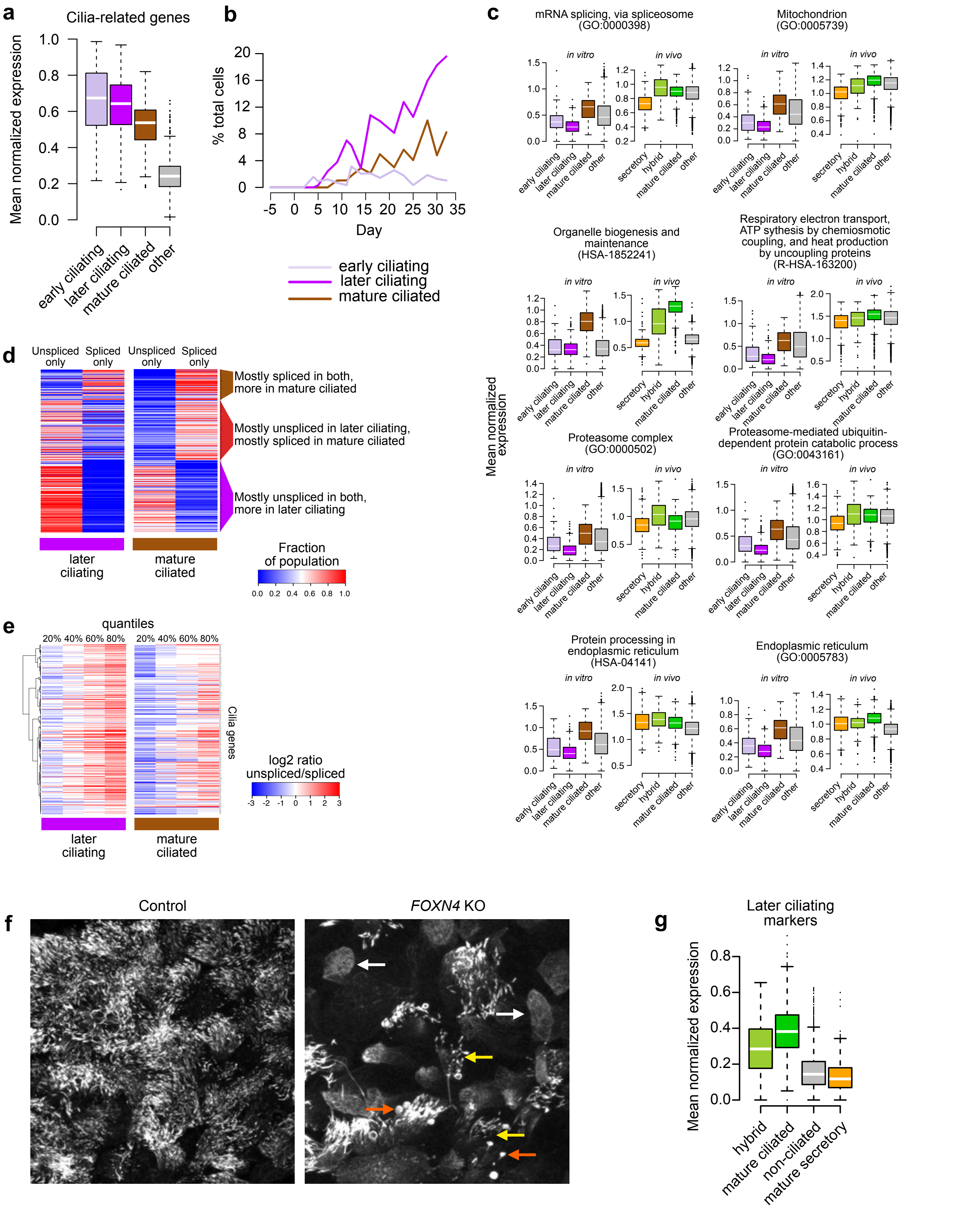

**Supplementary Figure S5 (with Figure 4): Relation of *in vitro* with *in vivo* ciliated cell states, splicing analysis and quantification of *FOXN4* KO phenotype**

- **a.** Mature ciliated cell genes common to the three *in vitro* ciliated populations distinguish these populations from the rest of the non-ciliated epithelium (“other”).
- **b.** Proportions of epithelial cells belonging to early ciliating, later ciliating and mature ciliated cell *in vitro* states across culture time points support the psuedotime trajectory in Supplementary Figure S4d.
- **c.** Enrichment analysis of DEGs upregulated in mature ciliated cells relative to later ciliating cells reveals that the mitochondrial maintenance and protein processing genes elevated in mature ciliated cells *in vitro* are also elevated in ciliating and ciliated cells *in vivo*. Box plots depict mean expression of genes enriched for the indicated term.
- **d.** Heat map illustrates differences in proportions for a given gene (one per row) that are spliced or unspliced between later ciliating and mature ciliated cells that exhibit non-zero expression for the gene. mRNA splice status was inferred using the Velocyto pipeline. Genes listed are cilia-related genes with non-zero expression (ignoring splicing) in at least 10% of cells for at least one of the later ciliating or mature ciliated cell populations.
- **e.** Ratio of unspliced/spliced RNA within later ciliating and mature ciliated cells suggests that specific ciliation program genes are poised (i.e., unspliced) in the later ciliating population, but active (i.e., spliced) in the mature ciliated cells. Quantiles of cells with both spliced and unspliced expression have a higher unspliced/spliced ratio across 243 cilia genes.
- **f.** ACT demonstrates example axonemal phenotypes used for immature ciliated cell quantification. FOXN4 KO cells displaying absent (white arrows), short or sparse (yellow arrows) or bulging (orange arrows) axonemes were classified as immature, while classic dense, well-formed axonemes (blue arrows) were classified as mature. Scale bar = 25 µm.
- **g.** Average expression of markers from the later ciliating *in vitro* state is higher in the hybrid secretory/ciliated state compared to secretory or other non-ciliated cells, reflecting its ciliating character. Other *in vitro* state marker expression profiles can be found in Figure 4e.

#### Supplementary Figure S6

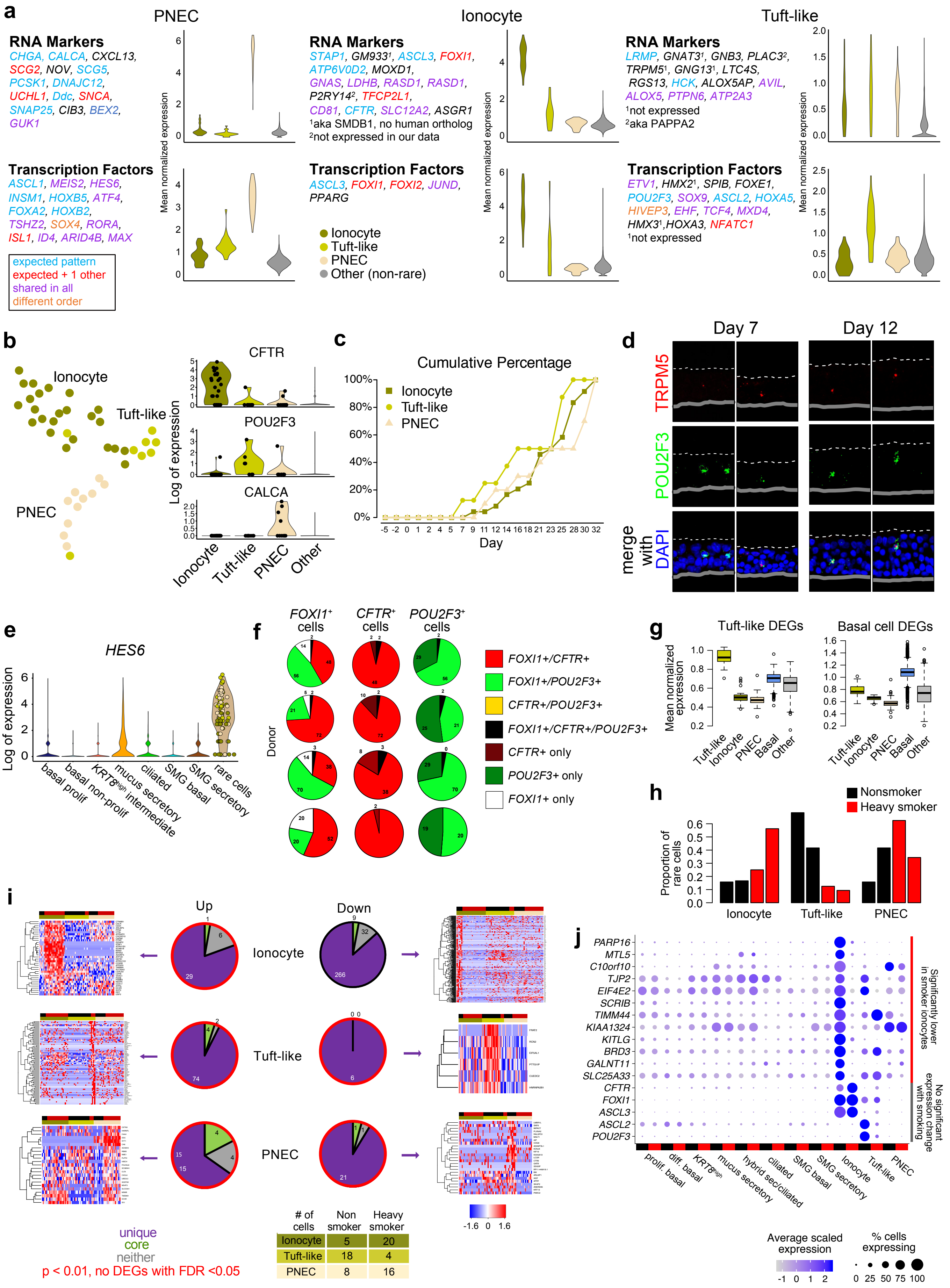

**Supplementary Figure S6 (with Figure 5): Human rare cell population comparisons with the literature, *in vitro* culture and smoking effects**

- **a.** Violin plots of average expression of recently published PNEC, ionocyte and tuft cell RNA markers and TFs[4]. Coloring legend is in the lower left, where blue text indicates genes that, in our dataset, follow the specific expression pattern presented by Montoro et al. (i.e. only present in the rare cell substate indicated), red text specifies genes that display the Montoro-predicted pattern as well as expression in 1 additional rare population, purple text indicates genes found to be expressed in all 3 rare populations in our dataset, and orange text denotes genes with expression in the Montoro-specified population, but higher expression in another rare cell population in our dataset.
- **b.** tSNE depicts SNN subclustering of rare cells found in ALI culture of human tracheal epithelial cells across all 20 time points in this study, and violin plots of ionocyte, tuft, and PNEC markers identify the three substates.
- **c.** Cumulative expression of ionocytes, PNECs and tuft-like cells captured by single cell sequencing at each time point in ALI culture.
- **d.** FISH localizes tuft-like cells to early time points in *in vitro* culture through co-localization of *POU2F3* (green) and *TRPM5* (red) mRNAs.
- **e.** Violin plots of *HES6* expression across rare cells and other populations *in vivo*. The color of the dots in the rare cell violin indicate to which rare cell population they belong, and correspond to the colors in **b**.
- **f.** Donor variation of *CFTR*, *FOXI1*, and *POU2F3* FISH quantification. Number of cells in indicated in each pie.
- **g.** *Left,* DEGs that distinguish Tuft-like cells from other rare cells have basal character compared to other rare cells and non-basal cells. The mean of all significant DEGs is plotted. *Right,* Tuft-like cells carry the most basal cell signature of all the rare cell types. The mean of the top 100 most upregulated genes in each of the differentiating and differentiating basal populations (compared to all other populations in the dataset) is shown.
- **h.** Proportion of rare cells belonging to each substate for each donor by scRNA-seq suggests a trending increase in ionocytes and decrease in tuft-like cells with heavy smoking.
- **i.** Rare cell smoking response trends: Pie charts depict the proportion and number of unique and core DEGs up (left) and down (right) in heavy smokers' rare cells. Pies encircled in red indicate genes that passed $p<0.01$ (no genes passed FDR adjustment in these comparisons). The pie encircled in black depicts genes that passed FDR<0.05 (discussed in the text with select genes in Figure 7G). Heat

maps beside each pie show the relative expression of all “unique” genes (purple pie slice) in comparison to the rest of the rare cell populations. Bars above the heat maps indicate donor (top), smoke status (middle; nonsmokers in black, heavy smokers in red), and rare cell type (bottom bar, colors correspond to the colors in **b**). The total number of cells in each smoke status are indicated in the table at the top center of the panel.

**j.** Ionocytes specifically lose a suite of metabolic genes in heavy smokers. Dot plots depict select genes of the 266 significantly uniquely downregulated genes in the ionocytes of heavy smokers, as well as known ionocyte and tuft genes for comparison.

**Supplementary Table S1: Demographic information for tracheal and lung tissue samples**

| Donor | Age | Sex | Total tobacco use (pack/day) | Years smoked | Years quit | Total pack years | Smoke status | *in vitro* too? |
|---|---|---|---|---|---|---|---|---|
| T84 | 22 | M | 1.5 packs/day | 2 years | 2 years | 3 | light | yes |
| T85 | 59 | M | 0.5 pack/day | 30 years | | 15 | heavy | yes |
| T89 | 10 | F | | | | 0 | never | yes |
| T120 | 57 | F | 3 pack/day | >30 years | | 90 | heavy | no |
| T121 | 23 | F | 0.5 pack/day | unknown | | <4 | light | no |
| T126 | 35 | F | | | | 0 | never | no |
| T137 | 27 | M | | | | 0 | never | no |

**Supplementary Table S2**: **All significant heavy smoking DEGs for each cluster or subcluster.** (SupplementaryTableS2_smokingDEGs.xlsx)

Seurat differential expression output tables of upregulated DEGs are listed first (in separate tabs) for each cluster/subcluster investigated, followed by tables for downregulated DEGs. In each table, “avg_logFC” = average log fold change, “p_val” = p-value, “p_adj_FDR” = FDR, “pct.1” = proportion of heavy smoker cells (for up genes) or nonsmoker cells (for down genes) expressing the gene, “pct.2” = proportion of nonsmoker cells (for up genes) or heavy smoker cells (for down genes) expressing the gene, “comparison” = the cluster/subcluster of cells involved and the direction of heavy smoking effect (up or down), and “DEG_status” = whether the DEG was “core” (observed in four or more populations, excluding the hybrid and rare cell groups), “unique” (observed in only that population), or “semi-unique” (not core or unique). The final two tabs, “core_up” and “core_down”, list “core” response DEGs that were up or downregulated, respectively.

#### Supplementary Table S3: Overlapping DEGs among rare cells

| Average expression<br>Ionocyte Tuft-like PNEC other | Number of significant genes | Key genes & pathways | Potential Regulators |
|---|---|---|---|
| 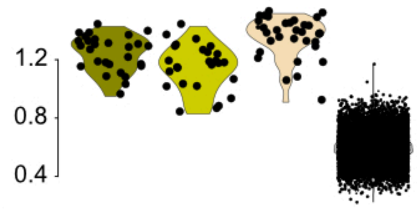 | 67 | *HEPACAM2 CLDN3 EPCAM BPHL RAB6A HACD3 SIGIRR BSG BSPRY SORL1*<br><br>transmembrane transport<br>secreted protein processing<br>cell adhesion | *HES6 ARID4B DHX36 GSE1 GTF3A H2AFY PBXIP1 STRAP FKBP1A* |
| 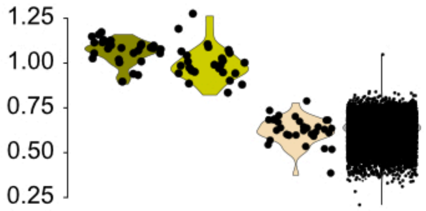 | 114 | *ILF2 KIT BIK CFL1 DBN1 MUC20 PTTG1IP SRGAP1 ANXA3 ANXA4 B3GNT2 BCL2 CXCR4 H3F3B HILPDA KRT18 KRT7 PRKRA RAMP2 SLIRP RBPJ ST14 DDAH2 DPP3 FARP1 NOSIP*<br><br>somatic stem cell population maintenance<br>immune response/detox<br>neuronal processes<br>MAPK/ERK | *TEAD2 ASCL3 CITED2 CXXC5 DMRT2 FOXI1 MTDH CREB1 FOXP1 KDM4A PBX1 ZFHX3 ZMIZ1* |
| 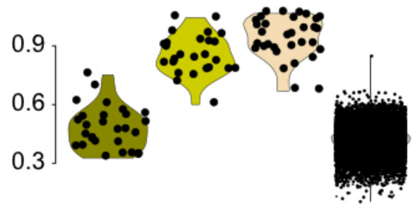 | 78 | *CRYM HYAL2 STMN1 MIF NPDC1 NRCAM SMS SPON2 CNPY2 DLL1 FBLIM1 GALNT7 GJC3 LPIN1 LRBA LSR MMP11 MTX2 NCAM1 RDH11 RPH3AL*<br><br>protein K48-linked ubiquitination<br>protein processing in the ER<br>microtubules in morphogenesis<br>neuronal processes | |
| 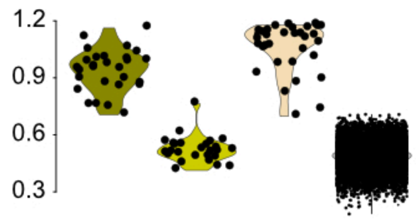 | 228 | *TMEM61 SEC11C ITPR2 NCALD GADD45G SCG2 HLA-A HLA-B HLA-C HLA-F PHB ANK2 QPCT NEURL1 CAPN2 RET TIMP2 NDFIP1 PHGR1 NOVA1 CD2AP PSAP TUSC3 IGFBP5 PTP4A3 ICA1 MAOB B2M VAMP*<br><br>insulin secretion<br>interferon signaling<br>protein processing in ER<br>calcium regulation | |

Violin plots of shared gene signature expression levels, genes and potential regulators across rare cell types implicate their relatedness.

#### Supplementary Table S4: Key resources for experimental methods

| REAGENT or RESOURCE | SOURCE | IDENTIFIER |
|---|---|---|
| **Antibodies** | | |
| chicken anti-KRT5 | BioLegend | 905901 |
| mouse anti-TP63 | Santa Cruz | sc8431 |
| rabbit anti-MKI67 | Abcam | ab16667 |
| rabbit anti-KRT8 | Abcam | ab81289 |
| mouse anti-MUC5AC | Invitrogen | MA-38223 |
| rabbit anti-MUC5B (H-300) | Santa Cruz | sc-20119 |
| mouse anti-KRT14 (LL002) | NeoMarkers | MS-115-P |
| rabbit anti-DMBT1 | Atlas Antibodies | HPA040778 |
| mouse anti-SCGB3A1 | Thermo | MA5-24147 |
| mouse anti-ACTA2 | Sigma | A2547 |
| goat anti-TSLP | BioLegend | 515901 |
| rabbit anti-FOXN4 | Thermo | PA5-62122 |
| mouse IgG2b anti-ac. alpha Tubulin | Sigma | T6793 |
| rabbit anti-DEUP1 (CCDC67) | Sigma | HPA010986 |
| mouse IgG1 anti-gamma Tubulin | Sigma | GTU88 |
| goat anti-rabbit IgG (H+L) | Invitrogen | A11008 |
| goat anti-mouse IgG2b | Invitrogen | A21242 |
| goat anti-mouse IgG1 | Invitrogen | A21125 |
| donkey anti-chicken IgY | Jackson ImmunoResearch | 703-545-155 |
| donkey anti-rabbit IgG (H+L) | Invitrogen | A21207, A31573 |
| donkey anti-goat IgG (H+L) | Invitrogen | A11058, A21447 |
| donkey anti-mouse IgG (H+L) | Invitrogen | A21203, A31571 |
| **Commercial Assays** | | |
| RNAScope Multiplex Fluorescent v2 Assay | Advanced Cell Diagnostics | 323100 |
| RNAScope probe Hs-SCGB1A1 | Advanced Cell Diagnostics | 469971 |
| RNAScope probe Hs-MUC5B | Advanced Cell Diagnostics | 449881 |
| RNAScope probe Hs-MUC5AC | Advanced Cell Diagnostics | 312891 |
| RNAScope probe Hs-IL33 | Advanced Cell Diagnostics | 400111 |
| RNAScope probe Hs-FOXJ1 | Advanced Cell Diagnostics | 430921 |
| RNAScope probe Hs-CFTR | Advanced Cell Diagnostics | 603291 |
| RNAScope probe Hs-CHGA | Advanced Cell Diagnostics | 311111 |
| RNAScope probe Hs-POU2F3 | Advanced Cell Diagnostics | 300031 |
| RNAScope probe Hs-ACTA2 | Advanced Cell Diagnostics | 311811 |
| RNAScope probe Hs-SOX9-No-XMm | Advanced Cell Diagnostics | 543021 |
| RNAScope probe Hs-FOXI1 | Advanced Cell Diagnostics | 476351 |
| RNAScope probe Hs-TRPM5 | Advanced Cell Diagnostics | 400891 |
| Alt-R HiFi Cas9 nuclease V3 | IDT | 1081061 |
| Alt-R CRISPR-Cas9 tracrRNA | IDT | 1072533 |
| Alt-R CRISPR-Cas9 Negative Control crRNA #1 | IDT | 1072544 |
| Alt-R FOXN4 crRNA 1 | IDT | AAGCTGTAGATCTCGCTCAC |
| Alt-R FOXN4 crRNA 2 | IDT | CACCGGAGTATGGCCAACCC |

| REAGENT or RESOURCE | SOURCE | IDENTIFIER |
|---|---|---|
| **Commercial Assays (cont)** | | |
| Amaxa Basic Nucleofector Kit, Primary Mammalian Epithelial Cells | Lonza | VPI-1005 |
| ICELL8 Chip Kit v2 | WaferGen | 430-000245 |
| Quick RNA MicroPrep Kit | Zymo | R1051 |
| Ion AmpliSeq Transcriptome Human Gene Expression Kit | Life Technologies | A26326 |
| **Chemicals** | | |
| DMEM-C | Corning | Corning-017 |
| DMEM-F | Fisher | SH3024301 |
| Ham's F-12 | Gibco | 11765-054 |
| FBS | LifeTechnologies | 10082-147 |
| L-glutamine | Gibco | 25030-081 |
| Penicillin/Streptomycin (P/S) | Gibco | 15140-122 |
| 100X Penicillin/Streptomycin/AmphtericinB (PSA) | Fisher | ICN1674049 |
| Hydrocortisone | Sigma | H-0888 |
| EGF | Gibco | 17005-042 |
| Cholera Toxin | Sigma | C-3012 |
| Insulin | Sigma | I9278 |
| Adenine | Sigma | A8626 |
| BEBM | Lonza | CC-3171 |
| DMEM | Gibco | 11885 |
| BEGM Brochial Epithelial Growth Medium BulletKit | Lonza | CC-4175 |
| PneumaCult ALI Medium Kit | Stemcell Technologies | 5001 |
| Rock Inhibitor (Y-27632 dihyrochloride) | APExBio | A3008 |
| Amphotericin B | sigma | A9528 |
| Fluconazole | Gallipot | NDC 51552-1031-2 |
| Gentamicin Reagent | Gibco | 15710-064 |
| Trypsin | Fisher | 25-053-CI |
| 1X DPBS | Fisher | 21-040-CV |
| 1X HBSS | Fisher | 21-021-CV |
| DNase | Sigma | DN25 |
| Accutase | Fisher | NC9839010 |
| HistoChoice | Sigma | H2779 |
| Antigen Unmasking Solution, Citrate based | Vector Labs | H-3300 |
| Protease from Streptomyces griseus | Sigma | P5147 |
| Collagen I, Rat | Corning | CB-40236 |
| PneumaCult-Ex Plus Medium | Stemcell Technologies | #05040 |
| **Other** | | |
| NIH 3T3 Fibroblasts (feeders) | ATCC | #CRL-1658 |
| 6.5mm transwell inserts, 0.4um | Corning | 3470 |
| Cell Strainer, 70um | Fisher | 08-771-2 |

**Supplementary Table S5**: **Specifications for donor alignment, clustering, and visualization for datasets and data subsets used in the study.**

| *Dataset* | *N top genes* | *N genes used* | *N donors* | *N CCA dims* | *Res* | *k* | *Optimizer* | *Plx* |
|---|---|---|---|---|---|---|---|---|
| All cells, *in vivo* | 8,000 | 5,009 | 5/7 | 25 | 0.3 | 30 | Louvain | 100 |
| All cells, *in vivo* (pre-culling)* | 2,500 | 1,790 | 5/7 | 29 | 0.4 | 30 | Louvain | 90 |
| All cells, *in vitro* | 10,000 | 9,542 | 2/3 | 20 | 0.4 | 15 | SLM | 80 |
| Rare cells, *in vivo* | 2,500 | 1,228 | 3/5** | 3 | 1.0 | 30 | Louvain | 19 |
| Rare cells, *in vitro* | 500 | 137 | 2/3 | 3 | 0.4 | 7 | SLM | 8 |
| SMG cells, *in vivo* (tSNE visualization only)† | 5,000 | 8,825 | 2/7 | 4 | NA | NA | NA | 60 |
| SMG basal cells, *in vivo* (subcluster assignment only)†† | 2,500 | 4,021 | 2/7 | 3 | 0.4 | 30 | Louvain | NA |
| Main basal cells, *in vivo* (subcluster assignment only)††† | 2,500 | 1,008 | 5/7 | 19 | 0.7 | 30 | Louvain | NA |
| Mucus secretory cells, *in vivo* | 8,000 | 3,573 | 5/7 | 12 | 0.4 | 30 | Louvain | 40 |
| KRT8$^{High}$ and mucus secretory cells, *in vivo* | 10,000 | 6,678 | 5/7 | 20 | 1.0 | 30 | Louvain | 130 |

*Footnotes*

* Includes non-epithelial populations; see Supplementary Figure S1a

** For dataset alignment and subclustering of the in vivo rare cell populations, we excluded two donors (T121 andT137) that only contained three rare cells each.

† Specifications for the tSNE visualization of all SMG populations (myoepithelial, SMG proliferating basal, SMG differentiating basal, and SMG secretory) shown in Supplementary Figure S6d.

†† Specifications for subclustering of the main SMG basal population (which include three differentiating basal and myoepithelial states) overlaid onto the tSNE plot in Supplementary Figure S6d.

††† Specifications for subclustering of the main proliferating and differentiating surface basal populations, which revealed the proliferating SMG basal population shown in Supplementary Figure S6d.

*Column descriptions*

*N top genes* = the number of top genes considered for each donor; *N genes used* = the total number of genes used for CCA; *N donors* = the number of donors (/out of all the donors available) needing the share a top gene for that gene to be used for CCA; *N CCA dims* = the number of CCA dimensions used for donor alignment; *Res*, *k*, and *Optimizer* = the resolution value, k value, and optimizer, respectively, used for SNN clustering; *Plx* = the perplexity used for tSNE visualization.

**Supplementary Table S6: Gene lists that accompany results presented in the main and Supplementary Figures.** (SupplementaryTableS6_GeneLists.xlsx)

Tables of DEGs are outputted from Seurat. In each table, “avg_logFC” = average log fold change (natural log), “p_val” = p-value, “p_adj_FDR” = FDR, “pct.1” = proportion of cells in the target population expressing the gene, “pct.2” = proportion of cells in the comparison population expressing the gene, and “comparison” = description of the comparison being made, usually with the target population listed first, and the comparison population listed second.

Tab 1. Major cluster DEGs (see Figure 1bc and Supplementary Figure S1c), where each cluster was compared against all the remaining clusters.

Tab 2. Markers for each of the major *in vivo* clusters. Differential expression results are given for the comparison yielding the highest p-value.

Tab 3. Upregulated and downregulated smoking response genes from Beane et al.[2] plotted in Figure 1i and Supplementary Figure S1f.

Tab 4. Genes associated with pseudotime in the mucus secretory cell trajectory in Figure 2ab (see also Supplementary Figure S2a). “module” identifies gene modules specified in the Figure (in addition to other “intermediate” modules not presented).

Tab 5. Significant enrichment results for the pseudotime-dependent genes in Tab 4.

Tab 6. Spearman partial correlation coefficients for genes significantly correlated with *MUC5AC*, *MUC5B*, or both (as indicated in the “which_mucin” column), accounting for smoking habit (heavy versus nonsmoking cells only). r = correlation coefficient, p = p-value, q = FDR. See Figure 2e for a scatter plot of these coefficients.

Tab 7. DEGs between two subclusters within the main mucus secretory cell population (mucus secretory vs. hybrid secretory/ciliated) (see Supplementary Figure S2b).

Tab 8. DEG table with 1) genes upregulated in surface (mucus) secretory cells compared to all non-secretory cells but *not* upregulated in SMG secretory cells compared to all non-secretory cells (“unique_to_surf.secretory”), 2) genes upregulated in SMG secretory cells compared to all non-secretory cells but *not* upregulated in surface secretory cells compared to all non-secretory cells (“unique_to_SMG.secretory”), and 3) genes upregulated in *both* surface and SMG secretory populations relative to non-secretory populations (“shared”). Differential expression results involving surface and SMG secretory populations are given in separate columns, as indicated by “surf.sec” and “SMG.sec” tags in the headings. See Figure 3a and Supplementary Figure S3a for heat maps of these genes.

Tab 9. DEGs among SMG basal cell states. Compare to the heat map of these genes in Supplementary Figure S3g.

Tab 10. Genes significantly associated with a pseudotime trajectory describing the transition from myoepithelial cells to SMG secretory cells through SMG basal cells (see Figure 3e and Supplementary Figure S3h). Genes were hierarchically clustered into 14 modules (A - N), which run from top to bottom in the heat map in Supplementary Figure S3h.

Tab 11. DEGs among cell clusters inferred for the *in vitro* time course dataset (see Supplementary Figure S4b) where each cluster was compared against all the remaining clusters.

Tab 12. Markers for each of the *in vitro* clusters.

Tab 13. Genes significantly associated with ciliated and secretory pseudotime lineages reconstructed for the *in vitro* time course dataset. Genes were clustered into five (ciliated) or four (secretory) modules that proceed from top to bottom in the heat maps in Supplementary Figure S4d.

Tab 14. Markers for each of the three *in vivo* rare cell populations. These include genes that, for each of the three groups, were both highly upregulated relative to other rare cells (FDR < 1e-5; see Tab 10) and highly upregulated relative to all other populations in the epithelium (FDR < 1e-5; see Tab 11). For each marker, differential expression results are shown from both comparisons.

Tab 15. DEGs for each *in vivo* rare cell population compared to other rare cells (see Figure 5b).

Tab 16. DEGs for each *in vivo* rare cell population compared to all other cells in the epithelium.

*References*

1 Fischer, A. J. *et al.* Differential Gene Expression in Human Conducting Airway Surface Epithelia and Submucosal Glands. *American Journal of Respiratory Cell Molecular Biology* **40**, 189-199 (2009).

2 Beane, J. *et al.* Reversible and permanent effects of tobacco smoke exposure on airway epithelial gene expression. *Genome Biology* **8** (2007).

3 Trapnell, C. *et al.* The dynamics and regulators of cell fate decisions are revealed by pseudotemporal ordering of single cells. *Nature Biotechnology* **32**, 381-U251, doi:10.1038/nbt.2859 (2014).

4 Montoro, D. T. *et al.* A revised airway epithelial hierarchy includes CFTR-expressing ionocytes. *Nature* **560**, 319-324 (2018).

5 Street, K. *et al.* Slingshot: cell lineage and pseudotime inference for single-cell transcriptomics. *BMC Genomics* **19**, 477-493 (2018).
